# Dopamine release and its control over early Pavlovian learning differs between the NAc core and medial NAc shell

**DOI:** 10.1101/2020.07.29.227876

**Authors:** Claire E. Stelly, Kasey S. Girven, Merridee J. Lefner, Kaitlyn M. Fonzi, Matthew J. Wanat

**Author notes:** **Corresponding Author:** Matthew J. Wanat Neurosciences Institute, Department of Biology, University of Texas at San Antonio, One UTSA Circle, San Antonio, TX 78249.

## Abstract

Dopamine neurons respond to cues to reflect the value of associated outcomes. These cue-evoked dopamine responses can encode the relative rate of reward in rats with extensive Pavlovian training. Specifically, a cue that always follows the previous reward by a short delay (high reward rate) evokes a larger dopamine response in the nucleus accumbens (NAc) core relative to a distinct cue that always follows the prior reward by a long delay (low reward rate). However, it was unclear if these reward rate dopamine signals are evident during early Pavlovian training sessions and across NAc subregions. To address this, we performed fast-scan cyclic voltammetry recordings of dopamine levels to track the pattern of cue- and reward-evoked dopamine signals in the NAc core and medial NAc shell. We identified regional differences in the progression of cue-evoked dopamine signals across training. However, the dopamine response to cues did not reflect the reward rate in either the NAc core or the medial NAc shell during early training sessions. Pharmacological experiments found that dopamine-sensitive conditioned responding emerged in the NAc core before the medial NAc shell. Together, these findings illustrate regional differences in NAc dopamine release and its control over behavior during early Pavlovian learning.

## Introduction

Learning to associate cues with rewarding outcomes is a fundamental process that underlies reward-driven behaviors. The mesolimbic dopamine system regulates behavioral responses toward reward-predictive cues [1,2]. In particular, dopamine neurons respond to cues to encode the value of the outcome in well-trained animals. These cue-evoked dopamine responses can convey prospective reward-related information, such as reward preference, reward size, and reward probability [3-7]. Our recent findings illustrate that cue-evoked dopamine release also signals retrospective reward-related information [8]. In this prior study, rats were trained on a Pavlovian task in which distinct cues signaled identical outcomes but differed in the time elapsed since the previous reward delivery. The Short Wait cue always followed the previous reward by a short delay (high reward rate) while the Long Wait cue always followed the previous reward by a long delay (low reward rate). We found a larger dopamine response to the Short Wait cue in the nucleus accumbens (NAc) core of rats with extensive Pavlovian training (> 24 sessions) [8]. While these results demonstrate that dopamine encodes the relative reward rate in well-trained animals, it was unclear how these signals develop during early learning and if they are uniformly broadcast throughout the medial NAc.

Dopamine’s role in reward learning has been primarily studied using Pavlovian tasks with a single cue-reward relationship [9,10]. In contrast, the difference in the dopamine response between cues has been primarily studied in well-trained animals [3,4,6,7]. As such, it is unclear how dopamine signals emerge when learning multiple cue-reward relationships simultaneously. Cue-evoked dopamine release could acquire value-related information through a multi-step process or a single-step process. For example, in a multi-step process cue-evoked dopamine release first signals an upcoming reward (independent of value) and over training conveys the relative difference in value between cues. Alternatively, in a single-step process the cue-evoked dopamine response will reflect differences in reward value as these signals first emerge during training.

In the current study we performed voltammetry recordings of NAc dopamine release during early Pavlovian learning to determine if value-related dopamine signals develop in a single- or multi-step process. Rats were trained on a Pavlovian task where distinct cues were associated with different reward rates [8]. The presence of reward rate encoding by cue-evoked dopamine release during early training sessions would suggest value-related dopamine signals emerge via a single-step process. In contrast, the absence of reward rate encoding during early training sessions would indicate that value-related signals emerge via a multi-step process. We performed dopamine recordings in the NAc core and the medial NAc shell, as cue-evoked dopamine responses are present in both NAc subregions [11-13]. Additionally, we pharmacologically inhibited dopamine receptors to determine if conditioned responding requires dopamine in the NAc core and/or medial NAc shell.

## Methods

### Subjects and surgery

The University of Texas at San Antonio Institutional Animal Care and Use Committee approved all procedures. Male CD IGS Sprague Dawley rats (Charles River Laboratories, RRID:RGD 734476) were pair-housed upon arrival, allowed *ad libitum* access to water and chow, and maintained on a 12 h light/dark cycle. Voltammetry electrodes were constructed by threading a 7 μm diameter carbon fiber through polyamide-coated silica tubing and sealed with epoxy [14]. The sensing end of the electrode was cut to a length of ~150 μm. Voltammetry electrodes were surgically implanted under isoflurane anesthesia in rats weighing 300 – 400 g. Electrodes were implanted bilaterally and targeted the NAc core (relative to bregma: 1.3 mm anterior; ± 1.3 mm lateral; 7.0 mm ventral) or the medial NAc shell (1.5 mm anterior; ± 0.6 mm lateral; 7.3 mm ventral). Rats were also implanted with a Ag/AgCl reference electrode that was placed under the skull at a convenient location. Bilateral stainless-steel guide cannulae (InVivo One) were implanted 1 mm dorsal to the NAc core or medial NAc shell. Following surgery, rats were single-housed for the duration of the experiment and allowed to recover for 1-3 weeks before behavioral procedures.

### Behavioral procedures

At ≥ 7 days post-surgery, rats were placed on mild dietary restriction to 90% of their free feeding weight, allowing for a weekly increase of 1.5%. Rats were handled regularly before behavioral testing commenced. All behavioral sessions occurred during the light cycle in operant boxes (Med Associates) with a grid floor, a house light, a recessed food tray equipped with an infrared beam-break detector, and auditory stimulus generators (white noise and 4.5 kHz tone). To familiarize the animals with the operant chamber and food retrieval from the tray, rats first received 1-2 magazine training sessions in which 20 unsignaled food pellets (45 mg, BioServ) were delivered at a 90 ± 15 s variable interval. Rats underwent 6 Pavlovian reward conditioning sessions as described previously [8]. Pavlovian sessions consisted of 50 trials where the termination of a 5 s audio CS (tone or white noise, counterbalanced across animals) resulted in the delivery of a single food pellet and illumination of the food port light for 4.5 s. Each session contained 25 Short Wait trials and 25 Long Wait trials delivered in pseudorandom order. The Short Wait CS was presented after a 20 ± 5 s ITI, and the Long Wait CS was presented after a 70 ± 5 s ITI. We monitored head entries into the food tray across training sessions. Conditioned responding was quantified as the change in the rate of head entries during the 5 s CS relative to the 5 s preceding the CS delivery [8,15]. We also quantified the latency to initiate a head entry during the CS.

### Pharmacology

Flupenthixol dihydrochloride (Tocris) was dissolved in sterile 0.9% NaCl. Rats received bilateral 0.5 μl microinjections of flupenthixol (10 μg/side) or vehicle into the nucleus accumbens core or shell at 0.25 μl/min. The injectors were removed 1 min after the infusion ended. Behavioral sessions commenced 30 min after the microinjections [15,16].

### Voltammetry recordings and analysis

Indwelling carbon fiber microelectrodes were connected to a head-mounted amplifier to monitor dopamine release in behaving rats using fast-scan cyclic voltammetry [8,14,15,17-19]. During voltammetric scans, the potential applied to the carbon fiber was ramped in a triangular waveform from −0.4 V (vs. Ag/AgCl) to +1.3 V and back at a rate of 400 V/s. Scans occurred at 10 Hz with the electrode potential held at −0.4 V between scans. Dopamine was chemically verified by obtaining high correlation of the cyclic voltammogram during a reward-related event with that of a dopamine standard (correlation coefficient *r*^2^ ≥ 0.75 by linear regression). Voltammetry data for a session were excluded from analysis if the detected voltammetry signal did not satisfy the chemical verification criteria [8,15,15]. Dopamine was isolated from the voltammetry signal using chemometric analysis [20] with a standard training set accounting for dopamine, pH and drift. The background for voltammetry recording analysis was set at 0.5 s before the CS onset. Trials were excluded if chemometric analysis failed to identify dopamine on > 25% of the data points. The change in dopamine concentration was estimated based on the average post-implantation electrode sensitivity (34 nA/μM)[14].

CS-evoked dopamine release was quantified as the mean dopamine response during the 5 s CS relative to the 5 s prior to the CS delivery [8,15]. The slope of the CS response was calculated as the difference in the mean dopamine levels during the peak (1.5 – 2 s) and the end of the CS (4.5 – 5 s) as a function of time. The US-evoked dopamine response was quantified as the mean dopamine response during the 2.5 s following the pellet delivery relative to the mean dopamine response during the 0.5 s preceding the pellet delivery. The cumulative dopamine response during trials (CS + US dopamine response) was calculated as the average dopamine signal over the 7 s following the CS onset relative to the 5 s prior to the CS delivery.

### Experimental design and statistical analysis

We performed statistical analyses in Graphpad Prism 8 and RStudio. All data are plotted as mean ± SEM. A mixed-effects model fit (restricted maximum likelihood method) was used to analyze effects on behavioral measures and dopamine responses. Data were analyzed in 5 trial bins for within-session analyses or averaged within session for full training analyses. The significance level was set to α = 0.05 for all tests. A repeated measures correlation was used to correlate dopamine signals and behavioral outcomes across sessions [21]. A list of statistical analyses is presented in **Supplementary Table 1**.

### Histology

Rats were anesthetized, electrically lesioned via the voltammetry electrodes, and perfused intracardially with 4% paraformaldehyde. Brains were extracted and post-fixed in the paraformaldehyde solution for a minimum of 24 hrs, then were transferred to 15 % and 30 % sucrose in phosphate-buffered saline. Tissue was cryosectioned and stained with cresyl violet. Implant locations were mapped to a standardized rat brain atlas [22].

## Results

### CS-evoked dopamine release in the NAc does not encode reward rate in early training sessions

Rats were trained on a Pavlovian delay conditioning task in which 5 s audio conditioned stimuli (CSs) signaled the delivery of a food reward (US). This task involved two trial types with distinct CSs. Both CSs resulted in the identical outcome (a single food pellet) but differed in the time elapsed since the previous reward (**Fig. 1A**). In Short Wait trials, the CS was presented 15 – 25 s following the previous reward delivery (high reward rate). In Long Wait trials, the CS was presented 65 – 75 s following the previous reward delivery (low reward rate). Training sessions consisted of 25 Short Wait trials and 25 Long Wait trials presented in a pseudorandom pattern so that the identity of the upcoming trial could not be predicted. Rats with extensive training on this task (> 24 sessions) exhibit a larger NAc core dopamine response to the Short Wait CS relative to the Long Wait CS [8]. While CS-evoked dopamine encodes reward rate in well-trained rats, it is unclear when this signal first emerges. Furthermore, it is unknown whether dopamine signaling uniformly reflects reward rate throughout the medial NAc.

**Figure 1.**
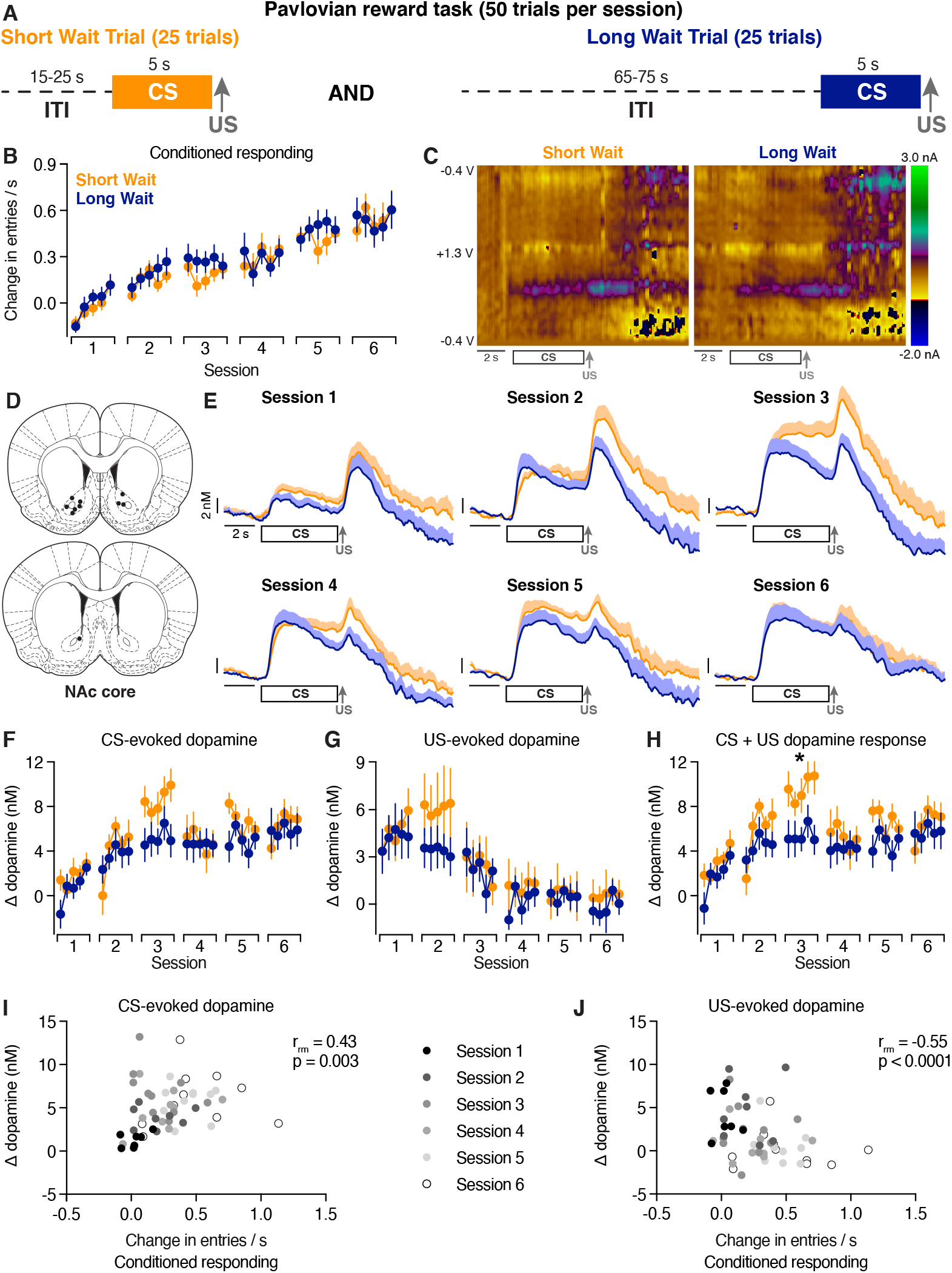
Dopamine release in the NAc core during early training sessions. (A) Pavlovian task. Short and Long Wait trials were presented in a pseudorandom pattern. (B) Conditioned responding across sessions in 5 trial bins. Conditioned responding was quantified as the change in the rate of head entries during the 5 s CS relative to the rate of head entries during the 5 s preceding the CS. (C) Representative two-dimensional pseudocolor plots of the resulting current from voltage sweeps (y-axis) as a function of time (x-axis) of voltammetry recordings in the NAc core. (D) Location of voltammetry electrodes. (E) Average dopamine signals across training sessions. (F) CS-evoked dopamine response across sessions in 5 trial bins. (G) US-evoked dopamine response across sessions in 5 trial bins. (H) Cumulative dopamine response during the CS and US across sessions in 5 trial bins. (I) Relationship between CS-evoked dopamine release and conditioned responding. (J) Relationship between US-evoked dopamine release and conditioned responding.

To address these questions, we performed voltammetry recordings of NAc dopamine levels in the NAc core and medial NAc shell during the first 6 Pavlovian training sessions. Conditioned responding in this task was quantified as the change in the rate of head entries during the 5 s CS relative to the rate of head entries during the 5 s preceding the CS [8]. Rats significantly increased conditioned responding to both CSs over all conditioning trials (two-way mixed effects analysis. trial effect *F*_(29,493)_=14.1, *p*<0.0001, n = 18 rats; **Fig. 1B** and **Supplementary Fig. 1**). The magnitude of conditioned responding did not differ between the Short and Long Wait CSs (reward rate effect *F*_(1,17)_=0.28, *p*=0.60), consistent with prior work [8]. Additionally, there was no difference in the latency to initiate a head entry between the trial types (**Supplementary Fig. 2**). We first examined the dopamine signals from electrodes in the NAc core (n = 10 electrodes; **Fig. 1C-E**). CS-evoked dopamine release was quantified as the average dopamine response during the 5 s CS relative to the 5 s prior to the CS delivery. The CS dopamine response increased over conditioning trials but did not differ between trial types (twoway mixed effects analysis. trial effect *F*_(5.0,44.9)_=5.54, *p*=0.0005; reward rate effect *F*_(1.0,9.0)_=1.71, *p=*0.22; **Fig. 1F**). These data illustrate that CS-evoked dopamine release in the NAc core does not reflect the reward rate during early training sessions.

We next examined US-evoked dopamine release by quantifying the average dopamine response during the 2.5 s following the pellet delivery relative to the average dopamine response during the 0.5 s preceding the pellet delivery. The US response decreased over all conditioning trials but did not differ between trial types (trial effect *F*_(3.6, 32.0)_=10.5, *p*< 0.0001; reward rate effect *F*_(1.0,9.0)_=0.208, *p*=0.66; **Fig. 1G**). We note that calculating US-evoked dopamine release in this manner obscures potential differences in dopamine levels immediately preceding the US delivery. As such, we also calculated the cumulative dopamine release during the CS and US by quantifying the average dopamine levels for the 7 s following the CS onset relative to the average dopamine levels during the 5 s preceding the CS (trial effect *F*_(2.1,12.6)_=5.7, *p*=0.02; reward rate effect *F*_(1.0, 6.0)_=2.2, *p*=0.19; **Fig. 1H**). A within-session analysis identified an elevated cumulative dopamine response during Short Wait trials in Session 3 (trial effect *F*_(2.7,24.0)_=1.2, *p*=0.33; reward rate effect *F*_(1.0,9.0)_=2.2, *p*=0.02; **Fig. 1H**). These results illustrate a transient difference in the total dopamine response between Short and Long Wait trials during early training sessions, which is distinct from the sustained difference in CS-evoked dopamine release in well-trained animals [8].

Recent optogenetic studies have produced conflicting results regarding dopamine’s involvement in conditioned responding, with some highlighting a role for dopamine during reward-predictive cues and others indicating a role for dopamine during the reward presentation [23-25]. Here, we examined how endogenous dopamine responses to the CS and US relate to conditioned responding across training sessions. Data were averaged between trial types as there were no differences in conditioned responding, CS-evoked dopamine release, or US-evoked dopamine release between Short and Long Wait trials. Conditioned responding was positively correlated with CS-evoked dopamine release and inversely related to US-evoked dopamine release in the NAc core (**Figs. 1I-J**).

We next examined dopamine signals from electrodes in the medial NAc shell (n = 14 electrodes; **Fig. 2A**). Similar to the NAc core, both the CS and US evoked time-locked phasic dopamine responses in the medial NAc shell (**Fig. 2B**). CS-evoked dopamine release increased over conditioning trials but did not differ between trial types (two-way mixed effects analysis: trial effect *F*_(5.0,65.1)_=4.93, *p*=0.0007; reward rate effect *F*_(1.0,13.0)_=0.00019, *p*=0.99; **Fig. 2C**). US-evoked dopamine release decreased over all conditioning trials but did not differ between trial types (trial effect *F*_(3.8, 49.0)_=9.83, *p*<0.0001; reward rate effect *F*_(1.0,13.0)_=2.33, *p*=0.15; **Fig. 2D**). Furthermore, the cumulative dopamine response across the CS and US was no different between Short and Long Wait trials (trial effect *F*_(5.1,6.0)_=4.0, *p*=0.0032; reward rate effect *F*_(1.0,13.0)_=0.06, *p*=0.81; **Fig. 2E**). When relating behavioral responding to dopamine signals we found that conditioned responding only correlated with US-evoked dopamine release in the medial NAc shell (**Figs. 2F-H**). Our results collectively demonstrate that the relative reward rate is not encoded by CS-evoked dopamine signals in either the NAc core or the medial NAc shell during early training sessions. In contrast, the CS-evoked dopamine response encodes the reward rate in rats with extensive Pavlovian training [8]. Together, these data indicate that value-related dopamine signals to reward-predictive cues emerge via a multi-step process.

**Figure 2.**
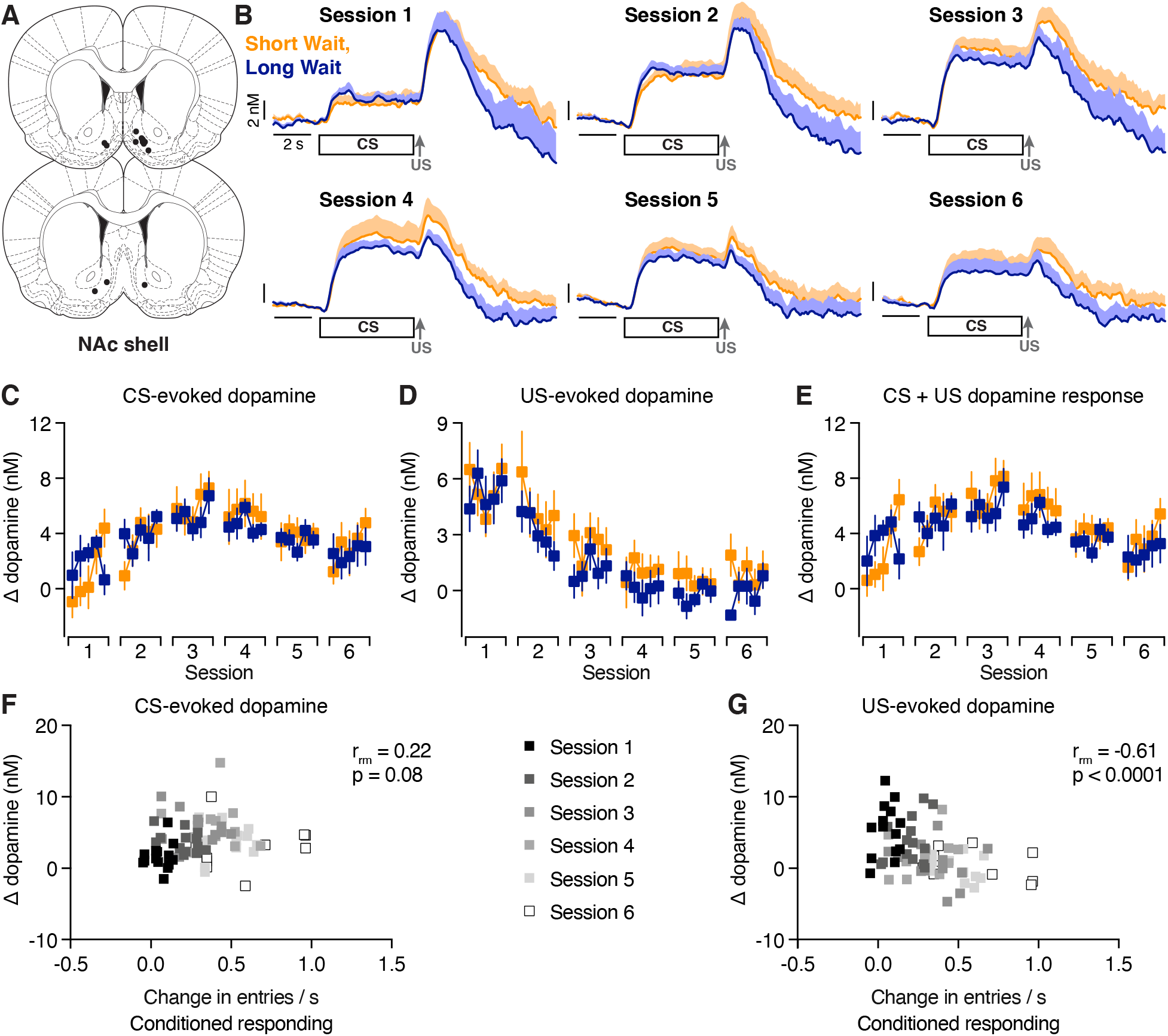
Dopamine release in the medial NAc shell during early training sessions. (A) Location of voltammetry electrodes. (B) Average dopamine signals across training sessions. (C) CS-evoked dopamine response across sessions in 5 trial bins. (D) US-evoked dopamine response across sessions in 5 trial bins. (E) Cumulative dopamine response during the CS and US across sessions in 5 trial bins. (F) Relationship between CS-evoked dopamine release and conditioned responding. (G) Relationship between US-evoked dopamine release and conditioned responding. (H) Relationship between the cumulative dopamine response and conditioned responding.

### Transient changes in the slope of the CS-evoked dopamine response

While there were no gross differences in CS-evoked dopamine release between Short and Long Wait trial types, the temporal dynamics of the response varied between trial types and across training sessions in the NAc core (**Fig. 1E**). To quantify these dynamics, we calculated the difference in dopamine levels from when average peak CS response occurred relative to the end of the CS presentation (**Fig. 3A**). In the NAc core, the slope of the CS response diverged between trial type in sessions 3 and 4 (two-way mixed effects analysis. reward rate effect in session 3 *F*_(1.0,9.0)_=13.8, *p*=0.0048; session 4 *F*_(1.0,8.0)_=10.3, *p*=0.013; **Fig. 3B**). This transient effect was no longer observed by session 5 (reward rate effect. *F*_(1.0,9.0)_=4.68, *p*=0.059). The difference in the slope of the CS dopamine response between trial types was not accompanied by a corresponding difference in conditioned responding or the latency to respond for either trial type (**Supplementary Fig. 3**). In contrast to the NAc core, the slope of the CS-evoked dopamine response did not differ between Short and Long Wait trial types in the medial NAc shell (reward rate effect for all trials. *F*_(1.0,13.0)_=0.0057, *p=*0.94; **Fig. 3C**). Together, these results highlight trialtype and region-specific changes in the dynamics of the CS-evoked dopamine response during early Pavlovian training sessions.

**Figure 3.**
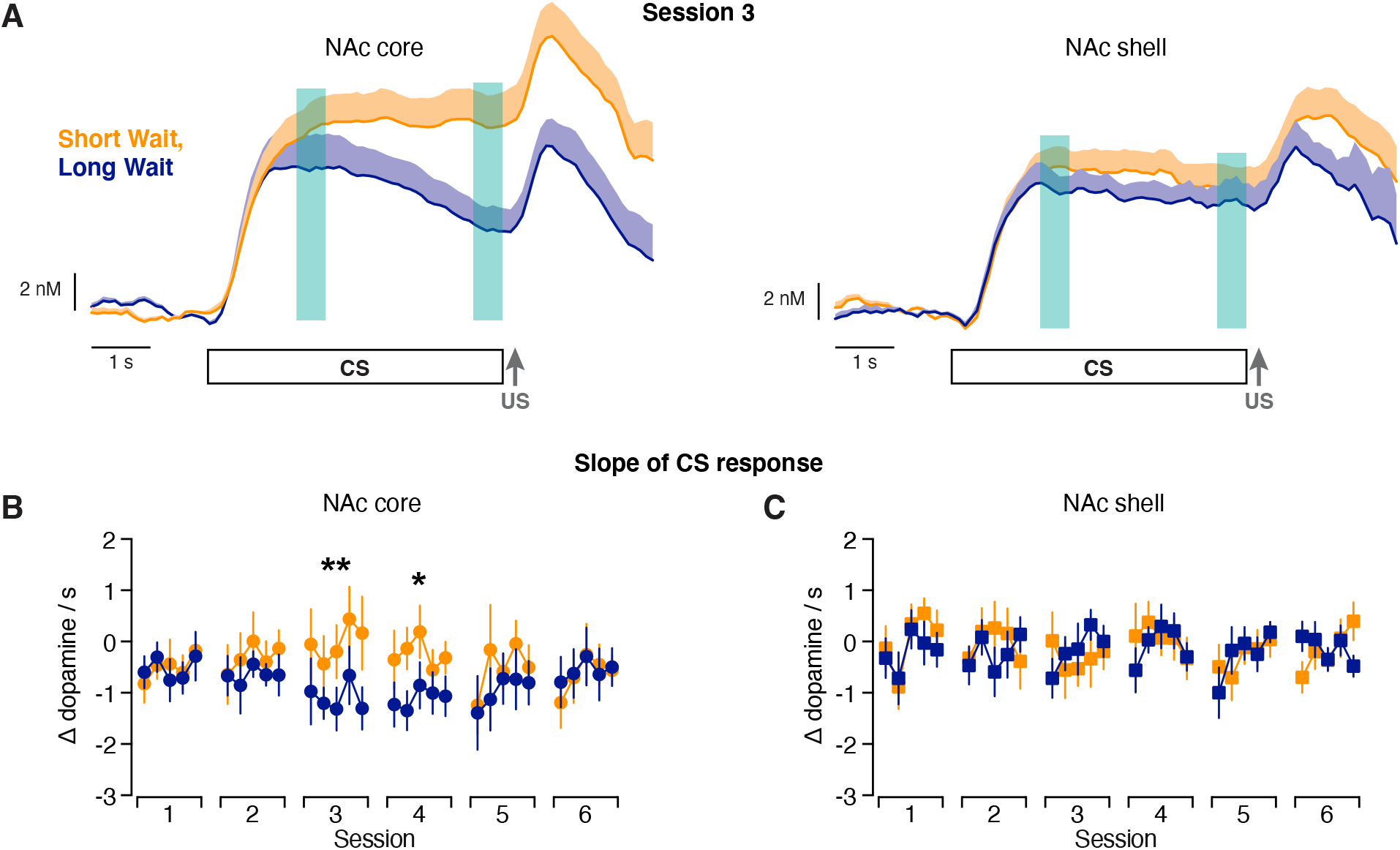
Differential dynamic of the CS dopamine response between trial types in the NAc core. (A) Slope of the CS dopamine response is calculated as the difference between the regions of interest denoted by the teal overlay. (B) Slope of the CS dopamine response in the NAc core. (C) Slope of the CS dopamine response in the medial NAc shell. * p < 0.05, ** p < 0.01 main effect of trial type.

### Regional differences in dopamine release

Prior studies have noted differences in dopamine release between the NAc core and NAc shell in reward-based tasks [11-13,26,27]. As such, we directly compared dopamine release between the NAc core and medial NAc shell during early Pavlovian training sessions. This analysis identified a significant interaction of brain region and training session on the CS-evoked dopamine response (three-way mixed effects analysis: region x session *F*_(5,84)_=2.93, *p=*0.017; region effect *F*_(1,84)_=2.08, *p*=0.154; session effect *F*_(2.0,43.3)_=12.15, *p*<0.0001; reward rate effect *F*_(1.0, 22.0)_=0.99, *p*=0.33; **Fig. 4A**). In contrast, US-evoked dopamine release did not differ between the NAc subregions (three-way mixed effects analysis: region x session *F*_(5,84)_=0.56, *p*=0.73; region effect *F*_(1,84)_=0.00396, *p*=0.950; session effect *F*_(1.60,35.2)_=28.3, *p*<0.0001; reward rate effect *F*_(1.0,22.0)_=1.41, *p*=0.25; **Fig. 4B**). There was a significant interaction of brain region and training session when examining the cumulative dopamine response during the trial (three-way mixed effects analysis. region x session *F*_(5,84)_=3.01, *p*=0.015; region effect *F*_(1,84)_=1.53, *p*=0.22; session effect *F*_(2.2,47.8)_=7.88, *p*=0.0008; reward rate effect *F*_(1.0,22.0)_=3.39, *p*=0.0079; **Fig. 4C**). The regional differences in dopamine release were driven by a smaller dopamine response in the medial NAc shell during later training sessions. Together, these results highlight the divergence of CS-evoked release and the cumulative trial dopamine response between the NAc core and medial NAc shell over the course of associative learning.

**Figure 4.**
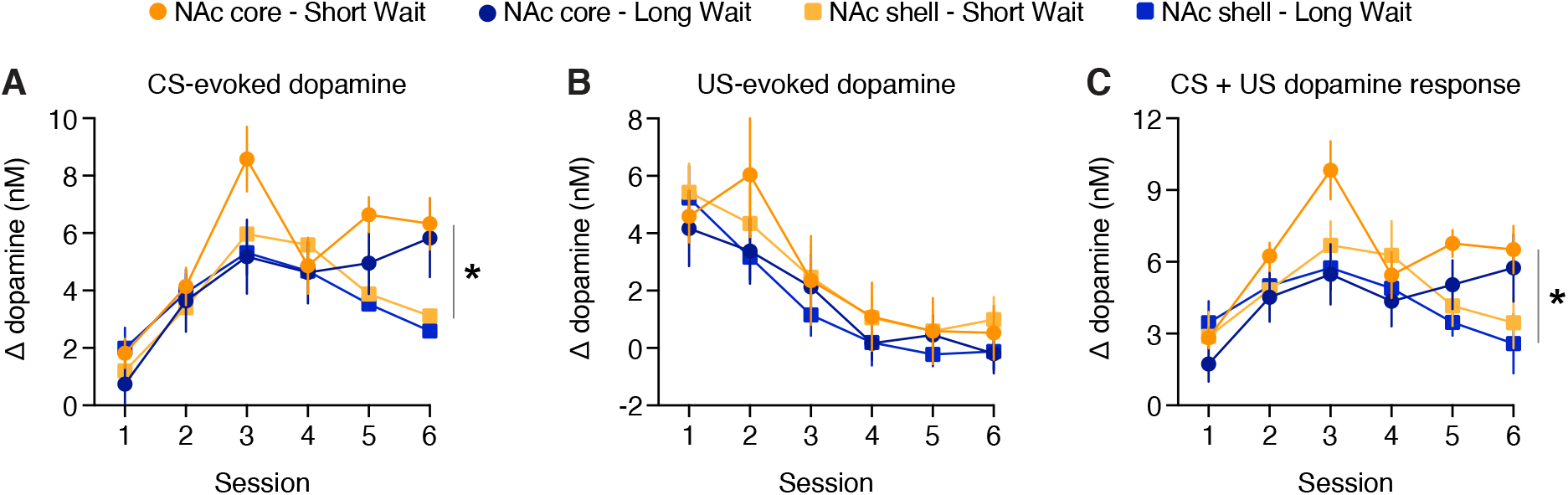
Comparing the dopamine response between the NAc core and medial NAc shell. (A) CS-evoked dopamine response. (B) US-evoked dopamine response. (C) Cumulative dopamine response during the CS and US. * p < 0.05 interaction effect of session x NAc core x medial NAc shell.

### Emergence of dopamine-sensitive conditioned responding in the NAc core and medial NAc shell

NAc dopamine signaling is necessary for conditioned behavioral responses to rewarding cues [16,28]. However, the differential development of the CS-evoked dopamine responses between the NAc core and medial NAc shell suggests that these regions may not contribute equally to behavior throughout training. To address this, rats were implanted with bilateral cannulae targeting the NAc core or the medial NAc shell for local pharmacological manipulations (**Fig. 5A**). The D1/D2 dopamine receptor antagonist flupenthixol (10 μg/side) or vehicle was infused 30 min before the first 5 sessions. Rats were trained without microinjections for an additional session to differentiate acute versus sustained behavioral effects of the drug treatment (**Fig. 5B**).Flupenthixol microinjections in the NAc core significantly disrupted conditioned responding (Drug effect *F*_(1,456)_=6.85; *p*=0.0091; Reward rate effect *F*_(1.0,19.0)_=1.85; *p*=0.19; Drug x Reward rate interaction *F*_(1,456)_=0.67; *p*=0.80; n = 10 vehicle, 11 flupenthixol; note that data are plotted separately by trial type for visual clarity in **Fig. 5C**). Within-session analyses identified lower levels of conditioned responding in flupenthixol-treated rats during the third and fifth training session (three-way mixed effects analysis. drug effect session 1 *F*_(1,76)_=1.97, *p*=0.17; session 2 *F*_(1,76)_=0.88, *p*=0.35; session 3 *F*_(1,76)_=4.16, *p*=0.045; session 4 *F*_(1,76)_=3.95, *p*=0.05; session 5 *F*_(1,76)_=10.3, *p*=0.0019). Furthermore, impairments in conditioned responding following flupenthixol treatment persisted during the sixth session in which no drug was administered (*F*_(1,76)_=11.5, *p*=0.0011). Flupenthixol application in the NAc core selectively reduced the number of CS-evoked head entries without altering head entries during the ITI or the latency to approach the food tray (**Supplementary Fig. 4**). Disruption of NAc core dopamine transmission therefore selectively impairs cue-driven appetitive behavior without altering motor function or response initiation.

**Figure 5.**
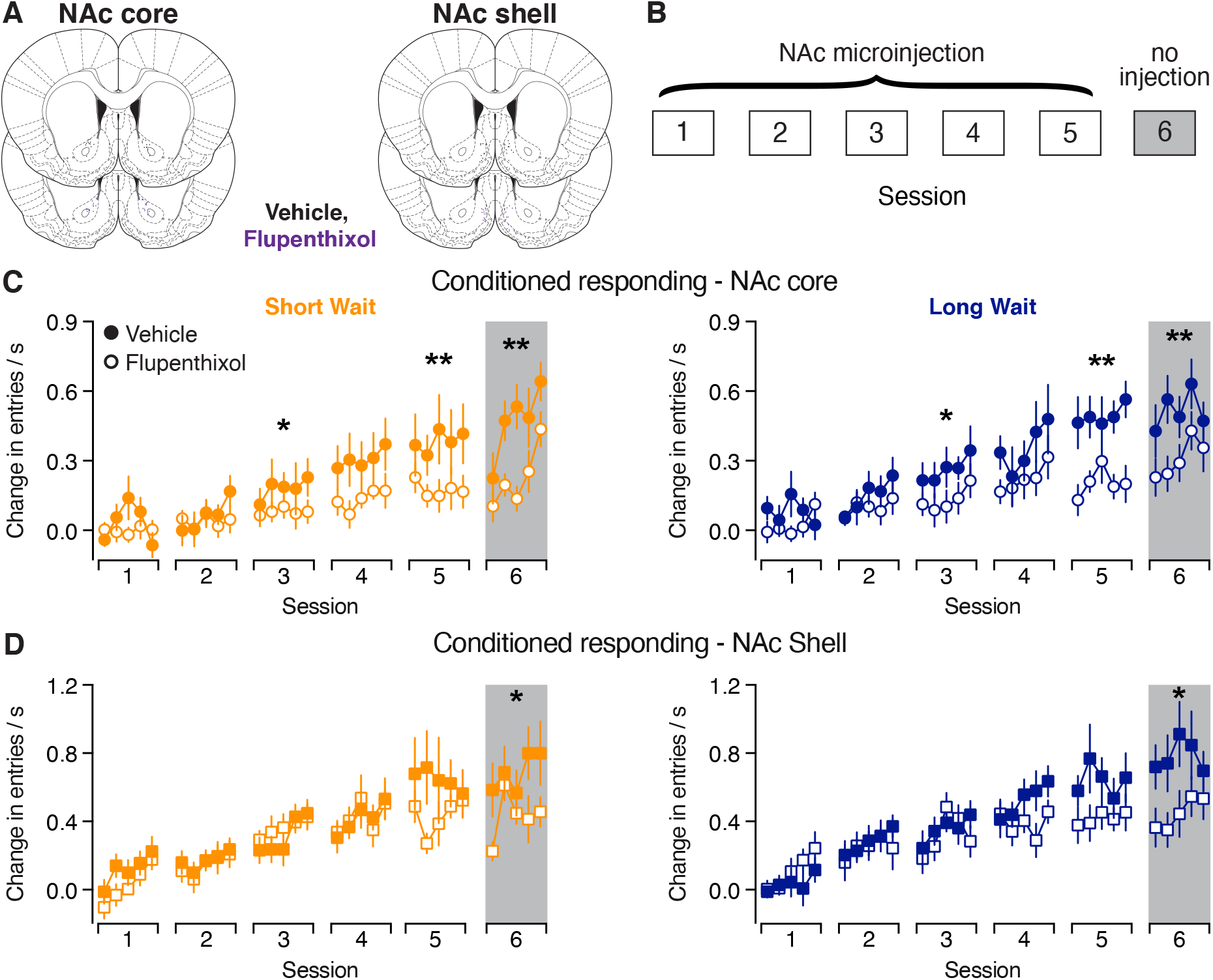
Emergence of dopamine-dependent conditioned responding in the NAc core and medial NAc shell. (A) Training paradigm. Flupenthixol or vehicle was infused into the NAc core or NAc shell before the first five training sessions. Training continued for an additional session without injections. (B) Location of the injector tips and infusion area. (C) Conditioned responding in rats receiving injections into the NAc core. (D) Conditioned responding in rats receiving injections into the NAc shell. Note that the data are plotted separately by trial type for visual clarity. * p < 0.05, ** p < 0.01 main effect of drug injection.

In rats with cannulae in the medial NAc shell, conditioned responding was not acutely affected by flupenthixol treatment during sessions 1-5 (Drug effect *F*_(1,408)_=1.16, *p*=0.28; Reward rate effect *F*_(1.0,17.0)_=0.19; *p*=0.67; Drug x Reward rate interaction *F*_(1,408)_=0.021; *p*=0.89; n = 10 vehicle, 9 flupenthixol; note that data are plotted separately by trial type for visual clarity in **Fig. 5D**). However, a behavioral deficit was evident on the sixth session in which no injection was administered (three-way mixed effects analysis: *F*_(1,68)_=5.06, *p*=0.028). Flupenthixol treatment in the NAc shell caused a subsequent non-significant reduction of cue-evoked head entries and significantly increased latency to approach the food tray during the sixth training session (**Supplementary Fig. 5**). This effect on response latency suggests that dopamine transmission in the NAc shell regulates the speed of response initiation in addition to the magnitude of the conditioned response. Collectively, these results demonstrate that dopamine signals in the NAc core and medial shell contribute to Pavlovian appetitive behavior at distinct phases of training.

## Discussion

Dopamine neurons respond to cues to convey reward-related information in Pavlovian tasks [3,7,8]. In particular, cue-evoked dopamine release in the NAc core signals the relative reward rate in rats with extensive Pavlovian training (>24 sessions) [8]. However, it was not known if reward rate dopamine signals rapidly emerged during early learning. To address this, we recorded dopamine release during the first 6 Pavlovian training sessions. Our data demonstrates that the reward rate is not reflected in cue-evoked dopamine responses in the NAc core or the medial NAc shell during early training. Coupled with our prior work, these findings suggest that NAc dopamine encodes value-related information via a sequential, multi-step process [8]. Specifically, cue-evoked dopamine release initially signals that a reward is forthcoming. Over training these cue-evoked dopamine responses then signal the relative value of the outcome.

The acquisition of conditioned responding depends on the temporal relationship between the cue, reward, and inter-trial interval (ITI). In particular, the ratio of the ITI duration relative to the cue duration can impact the rate of learning [29]. For example, an increase in the ITI (i.e. lower reward rate) facilitates acquisition in tasks involving a single cue-reward association [29-32]. However, we found no difference in the acquisition of conditioned responding between Short and Long Wait trials in our task. One potential explanation for this apparent discrepancy is that the difference in the ITI/cue ratio is not sufficiently large enough to observe differences in learning rates between the trial types. Alternatively, if rats cannot distinguish the difference in the ITI/cue ratios between the trial types in early training sessions, one would not anticipate a difference in the learning rate between Short and Long Wait trials.

Dopamine neurons projecting to the striatum are genetically and functionally diverse [23,33,33]. While rewards and reward-predictive cues elicit dopamine release in the NAc core and medial NAc shell, regional differences in the dynamics of the dopamine response are evident in some tasks [11-13,26,27]. Our data demonstrates dopamine release to cues initially increased in both regions, though there was a selective attenuation in the cue-evoked dopamine response in the medial NAc shell during later sessions. In contrast, there were no regional differences in the dopamine response to the reward delivery across sessions. Our results demonstrate that the increase in conditioned responding across sessions correlated with the decrease in reward-evoked dopamine release in the both the NAc core and medial NAc shell. In contrast, conditioned responding correlated with cue-evoked dopamine release in the NAc core, but not in the medial NAc shell. The selective relationship between conditioned responding and reward-evoked (and not cue-evoked) dopamine release in the medial NAc shell agrees with recent research demonstrating that the dopamine response to cues and rewards evolve independently of one another during early learning [35].

Although there was no difference in cue-evoked dopamine response between Short and Long Wait trials, we identified a transient trial type difference in the dynamics of the dopamine response in the NAc core. Specifically, the slope of the cue-evoked dopamine response diverged between trial types during session 3, which accounted for the greater cumulative dopamine release during Short Wait trials. These results indicate the presence of an additional factor that regulates dopamine transmission and potentially contributes to the initial discrimination between distinct cues [36-38]. We speculate that the slope of the dopamine response during the cue could be controlled by cholinergic signaling within the NAc. Striatal cholinergic neurons exhibit a reduction in firing to reward-predictive cues [39,40]. A decrease in cholinergic signaling can facilitate dopamine release evoked by high frequency stimulations [41,42]. The transient difference in the slope of the cue-evoked dopamine response between trial types could therefore arise from differences in striatal cholinergic signaling. Regardless, future studies will be needed to determine how striatal cholinergic neurons regulate the dynamics of dopamine release during early Pavlovian learning.

Conditioned responding can be modulated by the dopamine response to cues and rewards, though this can depend on task parameters and prior training [15,23-25]. We previously found that conditioned responding updates with cue-specific changes in dopamine release in well-trained animals. Rats trained to experience Short and Long Wait trials in separate sessions exhibited a selective elevation in dopamine release and conditioned responding to the Short Wait cue upon experiencing both trials together for the first time [8]. Based on these findings, we anticipated the emergence of cue-evoked dopamine release would parallel emergence of dopamine-sensitive conditioned responding during early training sessions. Our voltammetry recordings and dopamine receptor antagonist experiments instead illustrate that the increase in cue-evoked dopamine release precedes the emergence of dopamine-mediated conditioned responding. In the NAc core, flupenthixol treatment reduced conditioned responding starting in the third training session. These impairments persisted during the sixth training session with no drug treatment, which demonstrates NAc core dopamine is involved with Pavlovian learning, as reported previously [10,16]. In contrast, flupenthixol injections into the medial NAc shell failed to alter conditioned responding during the first five training sessions. However, this prior treatment with flupenthixol impaired conditioned responding during the sixth training session with no drug treatment. These results highlight a potential role for NAc shell dopamine in consolidation. In support, local injections of amphetamine into the NAc shell after Pavlovian training sessions resulted in elevated conditioned responding [43]. Future studies will be needed to identify the specific temporal window during the trial or after the session when dopamine signaling contributes to conditioned responding. Collectively, our results highlight regionspecific critical periods during training when dopamine signaling regulates conditioned responding.

Dopamine is thought to mediate distinct functions in the NAc core and NAc shell, with NAc core dopamine primarily involved with reward learning and NAc shell dopamine regulating learned behavioral actions [10,12,16,23]. However, it is important to note that these general roles may not be applicable to all behavioral tasks [44]. Indeed, our results highlight that dopamine in both the NAc core and medial NAc shell contribute to Pavlovian learning when cues convey distinct reward rates, albeit at different points during training. Future studies are needed to determine if dopamine in the NAc core and shell is similarly required for Pavlovian learning when cues signal differences in other reward-related parameters, such as reward size or probability. Research on dopamine’s role in behavior has largely focused on the ‘what’ and the ‘where’. what task elements increase dopamine release and where in the brain is dopamine released. Our results collectively highlight that it is also important to consider ‘when’ during training dopamine is capable of regulating behavioral actions.

## Author Contributions

CES, KSG, and MJW designed the experiments. CES, KSG, MJL, KMF, and MJW performed the experiments and analyzed the data. MJW and CES wrote the manuscript.

