## Supplementary Information for "Dopamine release and its control over early Pavlovian learning differs between the NAc core and medial NAc shell"

### Supplementary Figure 1

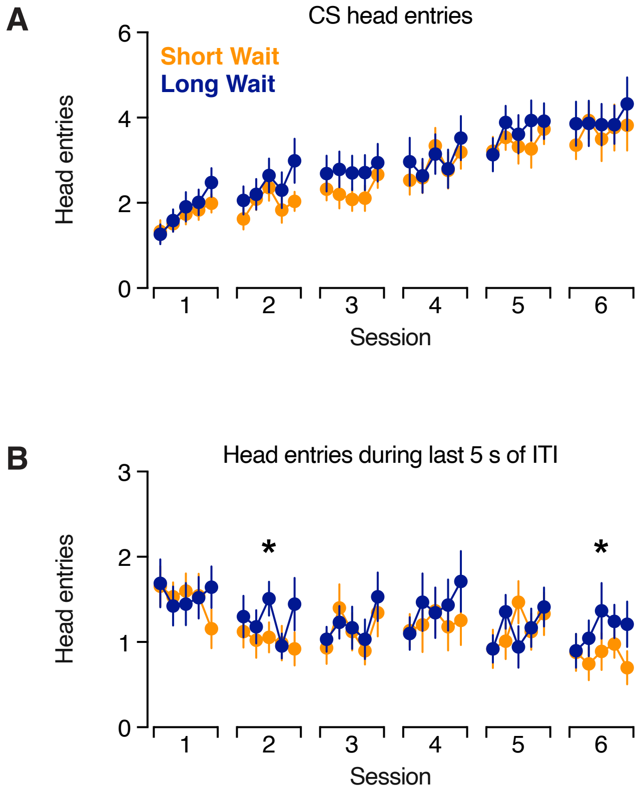


**Supplementary Figure 1**. (A) Head entries during five second cue during Short Wait (orange) and Long Wait (blue) trials (effect of trial type $F_{(1,17)}$= 1.17; $p$ = 0.295; effect of training $F_{(29,493)}$= 4.68; $p$ < 0.0001; interaction effect $F_{(29,493)}$= 0.747; $p$ = 0.829). (B) Head entries during the last five seconds of the inter-trial interval. Rats exhibit a higher level of anticipatory head entries on Long Wait trials, as reported previously [8] (effect of trial type $F_{(1,17)}$= 4.68; $p$ = 0.045; effect of training $F_{(29,493)}$= 1.81; $p$ = 0.0066; interaction effect $F_{(29,493)}$= 1.10; $p$ = 0.337). *$p$<0.05 main effect of trial type for within-session analysis.

### Supplementary Figure 2

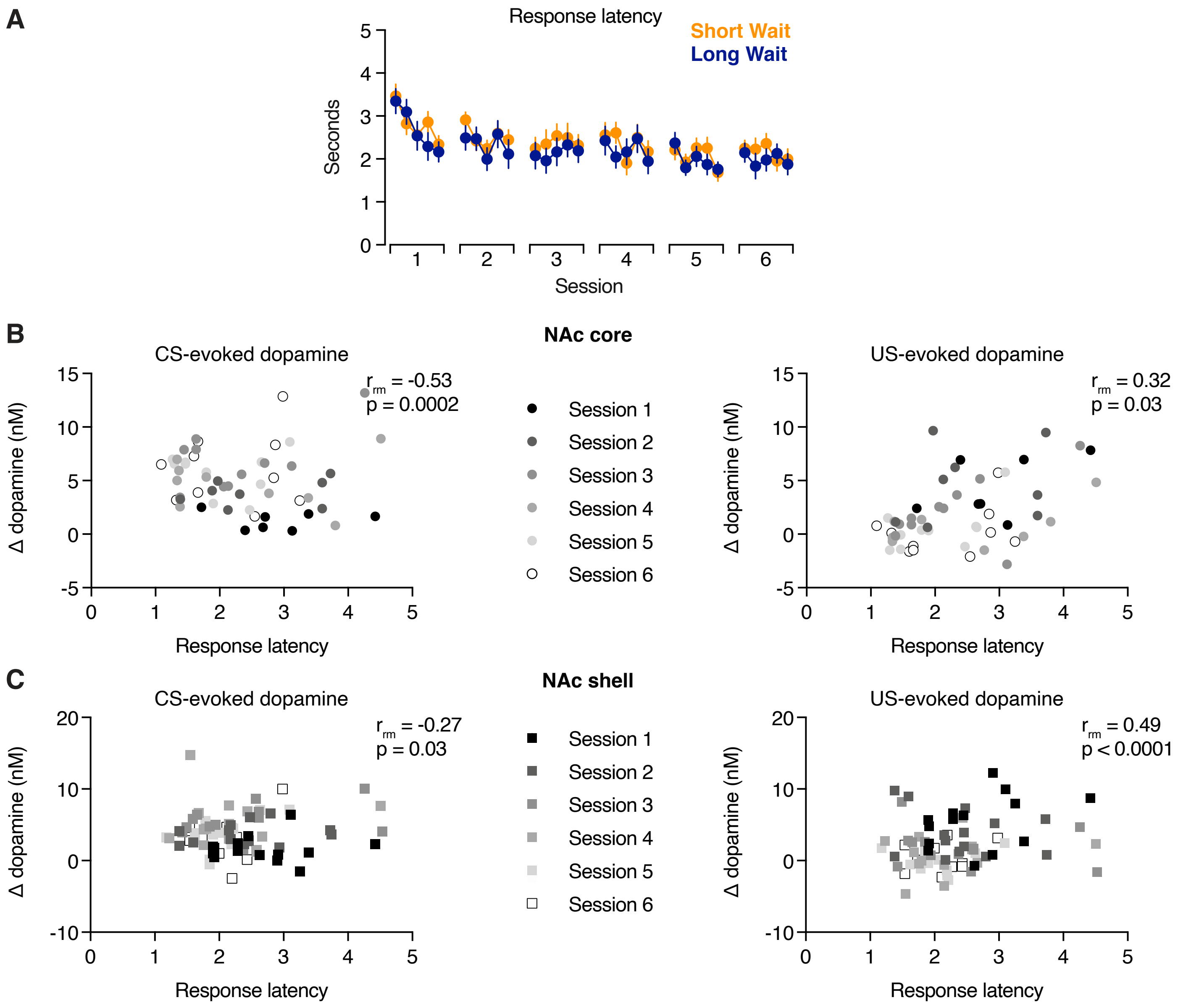


**Supplementary Figure 2**. Relationship between response latency and dopamine release. (A) Response latency across sessions in 5 trial bins. (B) Relationship between response latency and dopamine release in the NAc core evoked by the CS (left) and US (right). Both CS- and US-evoked dopamine respones correlate significantly with response latency. (C) Relationship between response latency and dopamine release in the NAc medial shell evoked by the CS (left) and US (right). Both CS- and US-evoked dopamine respones correlate significantly with response latency.

### Supplementary Figure 3

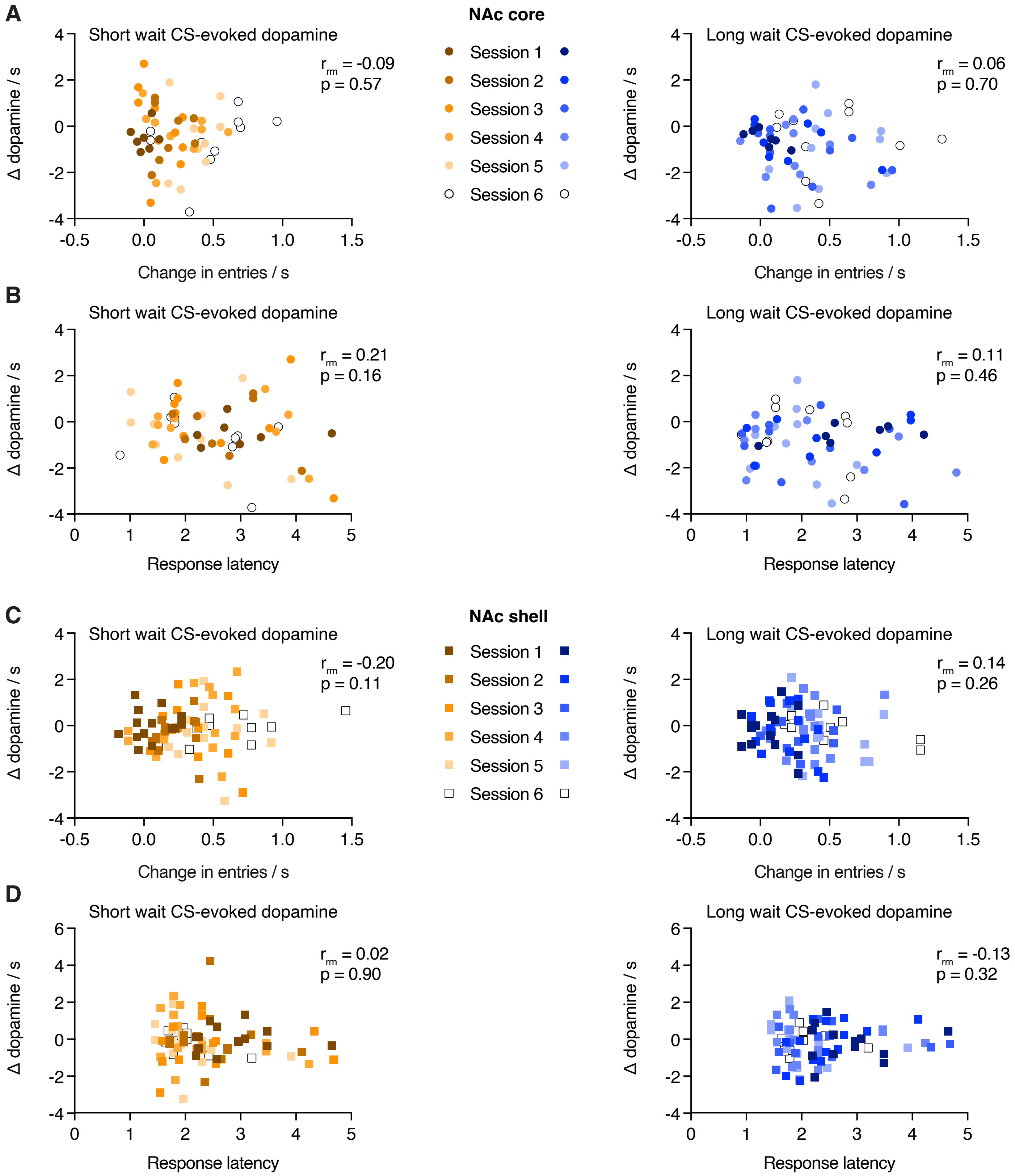


**Supplementary Figure 3**. Relationship between behavior and dopamine response dynamics. (A) Relationship between the slope of the CS-evoked dopamine response in the NAc core and conditioned responding in Short Wait (left, orange) and Long Wait (right, blue) trials. (B) Relationship between the slope of the CS-evoked dopamine response in the NAc core and response latency. (C) Relationship between the slope of the CS-evoked dopamine response in the NAc shell and conditioned responding. (D) Relationship between the slope of the CS-evoked dopamine response in the NAc shell and response latency.

### Supplementary Figure 4

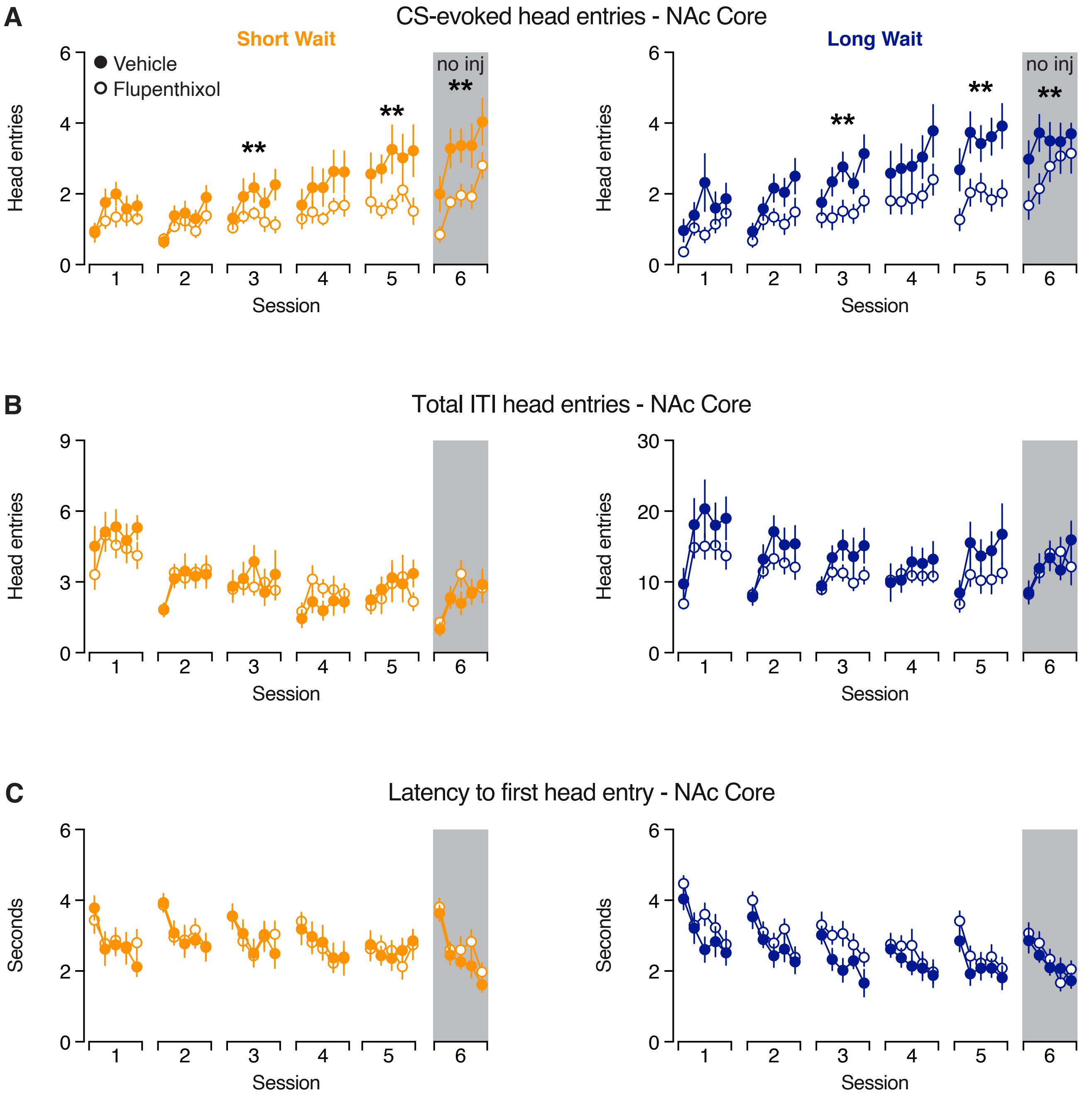


**Supplementary Figure 4**. Effect of flupenthixol treatment in the NAc core on cued and uncued responses. (A) Total head entries performed during the CS (Session 1-5: effect of trial type $F_{(1,19)}= 1.46$; $p = 0.241$; effect of flupenthixol $F_{(1,456)}= 9.12$; $p = 0.0027$; effect of training $F_{(7.9,150.9)}= 12.1$; $p < 0.0001$. Session 6: effect of trial type $F_{(1,19)}= 1.79$; $p = 0.196$; effect of flupenthixol $F_{(1,76)}= 7.93$; $p = 0.0062$; effect of training $F_{(3.0,57.1)}= 12.9$; $p < 0.0001$.) (B) Total head entries performed during the intertrial interval.(Session 1-5: effect of trial type $F_{(1,19)}= 0.035$; $p = 0.854$; effect of flupenthixol $F_{(1,456)}= 1.43$; $p = 0.233$; effect of training $F_{(8.9,168.6)}= 3.76$; $p = 0.0003$. Session 6: effect of trial type $F_{(1,19)}= 4.26$; $p = 0.053$; effect of flupenthixol $F_{(1,76)}= 0.00059$; $p = 0.981$; effect of training $F_{(3.0,56.1)}= 2.90$; $p = 0.044$.) (C) Latency to the first head entry from CS onset. (Session 1-5: effect of trial type $F_{(1,19)}= 0.602$; $p = 0.448$; effect of flupenthixol $F_{(1,456)}= 1.18$; $p = 0.279$; effect of training $F_{(10.2,193.1)}= 9.75$; $p < 0.0001$. Session 6: effect of trial type $F_{(1,19)}= 2.74$; $p = 0.114$; effect of flupenthixol $F_{(1,76)}= 1.76$; $p = 0.189$; effect of training $F_{(3.3,62.6)}= 19.7$; $p < 0.0001$.)   **$p < 0.01$ main effect of drug injection

### Supplementary Figure 5

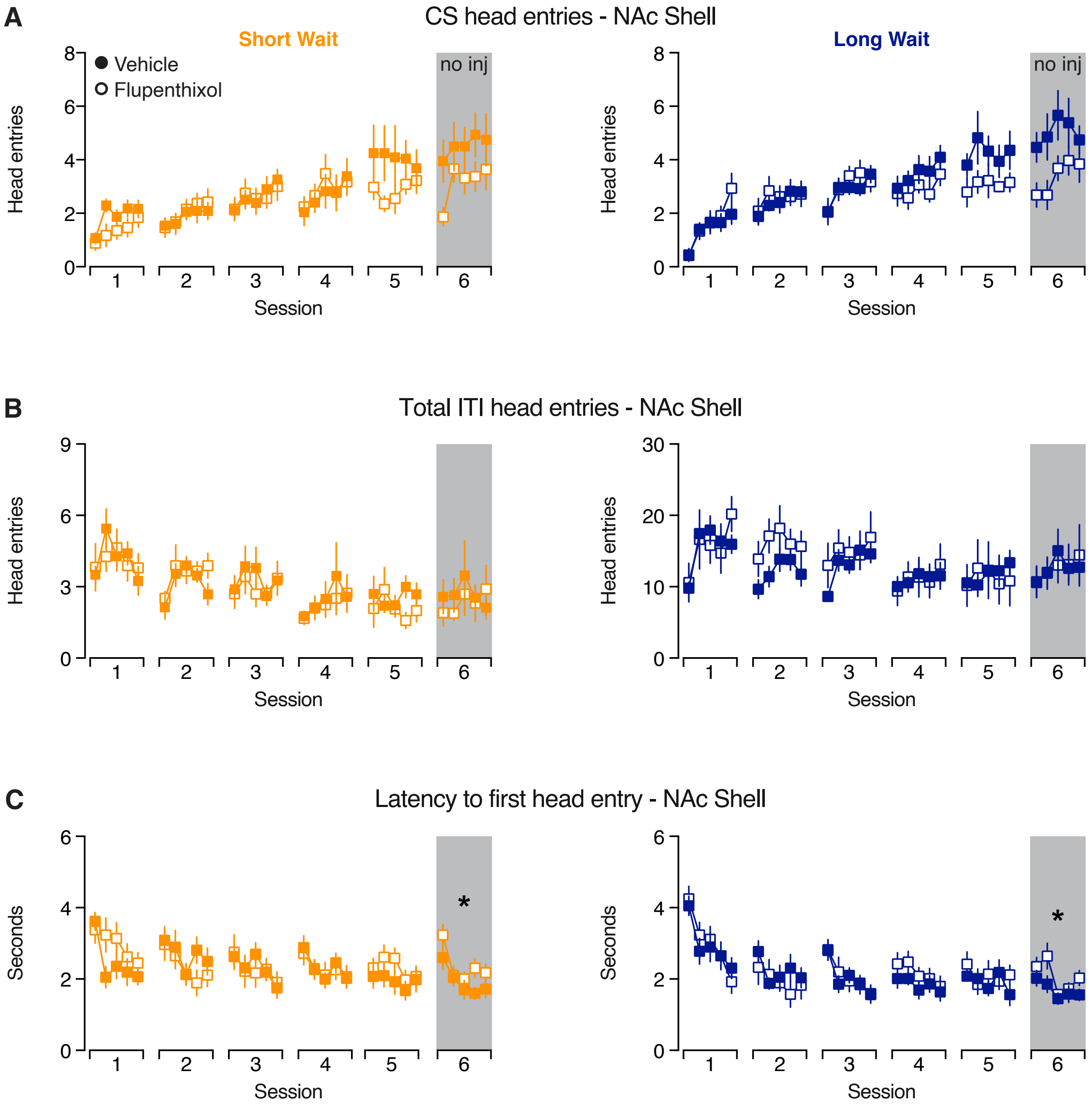


**Supplementary Figure 5**. Effect of flupenthixol treatment in the NAc shell on cued and uncued responses. (Session 1-5: effect of trial type $F_{(1,17)}$= 0.872; $p$ = 0.364; effect of flupenthixol $F_{(1,408)}$= 0.725; $p$ = 0.395; effect of training $F_{(7.3,123.5)}$= 13.1; $p$ < 0.0001. Session 6: effect of trial type $F_{(1,17)}$= 0.644; $p$ = 0.434; effect of flupenthixol $F_{(1,68)}$= 3.97; $p$ = 0.050; effect of training $F_{(3.3,55.5)}$= 4.13; $p$ = 0.0086.) (A) Total head entries performed during the CS. (B) Total head entries performed during the intertrial interval. (Session 1-5: effect of trial type $F_{(1,17)}$= 8.16; $p$ = 0.011; effect of flupenthixol $F_{(1,408)}$= 0.035; $p$ = 0.851; effect of training $F_{(9.7,164.5)}$= 2.86; $p$ = 0.003. Session 6: effect of trial type $F_{(1,17)}$= 0.677; $p$ = 0.422; effect of flupenthixol $F_{(1,68)}$= 0.005; $p$ = 0.941; effect of training $F_{(2.9,50.1)}$= 3.05; $p$ = 0.038.) (C) Latency to the first head entry from CS onset. (Session 1-5: effect of trial type $F_{(1,17)}$= 0.998; $p$ = 0.332; effect of flupenthixol $F_{(1,408)}$= 0.196; $p$ = 0.658; effect of training $F_{(8.6,145.7)}$= 8.07; $p$ < 0.0001. Session 6: effect of trial type $F_{(1,17)}$= 2.80; $p$ = 0.113; effect of flupenthixol $F_{(1,68)}$= 6.23; $p$ = 0.015; effect of training $F_{(3.5,58.7)}$= 12.3; $p$ < 0.0001.) *p<.05 main effect of drug injection

**Supplementary Table: Summary of Statistical Tests**

| **Figure 1** | | | |
|---|---|---|---|
| **Panel B – Conditioned Responding Behavior** | | | |
| All trials<br>Two-way mixed-effects model | Trial<br>$F_{(29,493)}$=14.1; **$p$<0.0001** | Reward rate<br>$F_{(1,17)}$=0.281; $p$=0.603 | Trial x reward rate<br>$F_{(29,493)}$=0.815; $p$=0.743 |
| Session 1<br>Two-way mixed-effects model | Trial<br>$F_{(4,68)}$=8.16; **$p$<0.0001** | Reward rate<br>$F_{(1,17)}$=0.345; $p$=0.565 | Trial x reward rate<br>$F_{(4,68)}$=0.345; $p$=0.847 |
| Session 2<br>Two-way mixed-effects model | Trial<br>$F_{(4,68)}$=3.56; **$p$=0.011** | Reward rate<br>$F_{(1,17)}$=0.357; $p$=0.558 | Trial x reward rate<br>$F_{(4,68)}$=1.07; $p$=0.379 |
| Session 3<br>Two-way mixed-effects model | Trial<br>$F_{(4,68)}$=0.818; $p$=0.518 | Reward rate<br>$F_{(1,17)}$=1.27; $p$=0.275 | Trial x reward rate<br>$F_{(4,68)}$=1.03; $p$=0.398 |
| Session 4<br>Two-way mixed-effects model | Trial<br>$F_{(4,68)}$=1.14; $p$=0.346 | Reward rate<br>$F_{(1,17)}$=0.0260; $p$=0.874 | Trial x reward rate<br>$F_{(4,68)}$=0.782; $p$=0.541 |
| Session 5<br>Two-way mixed-effects model | Trial<br>$F_{(4,68)}$=0.370; $p$=0.829 | Reward rate<br>$F_{(1,17)}$=0.567; $p$=0.462 | Trial x reward rate<br>$F_{(4,68)}$=1.83; $p$=0.132 |
| Session 6<br>Two-way mixed-effects model | Trial<br>$F_{(4,68)}$=1.44; $p$=0.231 | Reward rate<br>$F_{(1,17)}$=0.0122; $p$=0.913 | Trial x reward rate<br>$F_{(4,68)}$=0.711; $p$=0.587 |
| **Panel F – CS-evoked Dopamine** | | | |
| All trials<br>Two-way mixed-effects model | Trial<br>$F_{(5.0,44.9)}$=5.54; **$p$=0.0005** | Reward rate<br>$F_{(1.0,9.0)}$=1.71; $p$=0.224 | Trial x reward rate<br>$F_{(5.3,36.3)}$=1.58; $p$=0.188 |
| Session 1<br>Two-way mixed-effects model | Trial<br>$F_{(2.1,12.7)}$=4.10; **$p$=0.040** | Reward rate<br>$F_{(1.0,6.0)}$=1.18; $p$=0.319 | Trial x reward rate<br>$F_{(2.4,14.4)}$=1.52; $p$=0.253 |
| Session 2<br>Two-way mixed-effects model | Trial<br>$F_{(2.7,18.9)}$=5.99; **$p$=0.0058** | Reward rate<br>$F_{(1.0,7.0)}$=0.161; $p$=0.700 | Trial x reward rate<br>$F_{(2.0,13.8)}$=1.54; $p$=0.250 |
| Session 3<br>Two-way mixed-effects model | Trial<br>$F_{(2.9,26.3)}$=1.43; $p$=0.256 | Reward rate<br>$F_{(1.0,9.0)}$=4.23; $p$=0.0698 | Trial x reward rate<br>$F_{(3.0,26.6)}$=1.69; $p$=0.194 |
| Session 4<br>Two-way mixed-effects model | Trial<br>$F_{(1.8,14.5)}$=0.188; $p$=0.809 | Reward rate<br>$F_{(1.0,8.0)}$=0.0339; $p$=0.858 | Trial x reward rate<br>$F_{(1.9,15.1)}$=0.525; $p$=0.592 |
| Session 5<br>Two-way mixed-effects model | Trial<br>$F_{(2.6, 23.2)}$=2.04; $p$=0.144 | Reward rate<br>$F_{(1.0,9.0)}$=1.95; $p$=0.196 | Trial x reward rate<br>$F_{(2.4,20.8)}$=3.68; **$p$=0.0363** |
| Session 6<br>Two-way mixed-effects model | Trial<br>$F_{(2.0,17.6)}$=1.70; $p$=0.212 | Reward rate<br>$F_{(1.0,9.0)}$=0.289; $p$=0.604 | Trial x reward rate<br>$F_{(1.9,16.7)}$=2.01; $p$=0.167 |
| **Panel G – US-evoked Dopamine** | | | |
| All trials<br>Two-way mixed-effects model | Trial<br>$F_{(3.6,32.0)}$=10.5; **$p$<0.0001** | Reward rate<br>$F_{(1.0,9.0)}$=0.208; $p$=0.659 | Trial x reward rate<br>$F_{(3.1,21.4)}$=0.638; $p$=0.604 |
| Session 1<br>Two-way mixed-effects model | Trial<br>$F_{(3.0,18.1)}$=1.88; $p$=0.169 | Reward rate<br>$F_{(1.0,6.0)}$=0.299; $p$=0.604 | Trial x reward rate<br>$F_{(2.6,15.4)}$=0.850; $p$=0.472 |
| Session 2<br>Two-way mixed-effects model | Trial<br>$F_{(2.8,19.3)}$=0.0950; $p$=0.953 | Reward rate<br>$F_{(1.0,7.0)}$=1.26; $p$=0.299 | Trial x reward rate<br>$F_{(1.3,9.1)}$=0.352; $p$=0.624 |
| Session 3<br>Two-way mixed-effects model | Trial<br>$F_{(2.4,21.5)}$=1.98; $p$=0.156 | Reward rate<br>$F_{(1.0,9.0)}$=0.0135; $p$=0.910 | Trial x reward rate<br>$F_{(2.5,22.3)}$=1.61; $p$=0.219 |
| Session 4<br>Two-way mixed-effects model | Trial<br>$F_{(2.7,21.2)}$=1.19; $p$=0.335 | Reward rate<br>$F_{(1.0,8.0)}$=0.447; $p$=0.523 | Trial x reward rate<br>$F_{(3.0,23.7)}$=1.00; $p$=0.409 |
| Session 5<br>Two-way mixed-effects model | Trial<br>$F_{(2.3, 20.8)}$=0.190; $p$=0.857 | Reward rate<br>$F_{(1.0,9.0)}$=0.0106; $p$=0.920 | Trial x reward rate<br>$F_{(2.3,20.7)}$=0.339; $p$=0.746 |
| Session 6<br>Two-way mixed-effects model | Trial<br>$F_{(1.4,12.9)}$=0.622; $p$=0.501 | Reward rate<br>$F_{(1.0,9.0)}$=0.889; $p$=0.371 | Trial x reward rate<br>$F_{(2.7,24.7)}$=0.723; $p$=0.536 |
| **Panel H – CS+US Dopamine Response** | | | |
| All trials<br>Two-way mixed-effects model | Trial<br>$F_{(5.1,45.8)}$=1.23; **$p$=.0029** | Reward rate<br>$F_{(1.0,9.0)}$=5.90; **$p$=0.0380** | Trial x reward rate<br>$F_{(5.6,38.8)}$=1.86; $p$=0.117 |
| Session 1<br>Two-way mixed-effects model | Trial<br>$F_{(2.1,12.6)}$=5.71; **$p$=.0162** | Reward rate<br>$F_{(1.0,6.0)}$=2.17; $p$=0.191 | Trial x reward rate<br>$F_{(2.3,14.0)}$=1.01; $p$=0.401 |
| Session 2<br>Two-way mixed-effects model | Trial<br>$F_{(2.7,18.6)}$=6.11; **$p$=0.0056** | Reward rate<br>$F_{(1.0,7.0)}$=1.79; $p$=0.223 | Trial x reward rate<br>$F_{(2.0,14.2)}$=1.74; $p$=0.212 |
| Session 3<br>Two-way mixed-effects model | Trial<br>$F_{(2.7,24.0)}$=1.19; $p$=0.333 | Reward rate<br>$F_{(1.0,9.0)}$=8.79; **$p$=0.0159** | Trial x reward rate<br>$F_{(2.9,25.7)}$=1.48; $p$=0.243 |
| Session 4<br>Two-way mixed-effects model | Trial<br>$F_{(1.9,15.4)}$=0.185; $p$=0.826 | Reward rate<br>$F_{(1.0,8.0)}$=0.992; $p$=0.349 | Trial x reward rate<br>$F_{(2.1,17.1)}$=0.664; $p$=0.537 |
| Session 5<br>Two-way mixed-effects model | Trial<br>$F_{(3.5,31.7)}$=1.57; $p$=0.212 | Reward rate<br>$F_{(1.0,9.0)}$=3.68; $p$=0.0874 | Trial x reward rate<br>$F_{(2.4,21.7)}$=2.60; $p$=0.0889 |

| | | | |
|---|---|---|---|
| Session 6<br>Two-way mixed-effects model | Trial<br>$F_{(2.0,17.6)}$=2.54; $p$=0.108 | Reward rate<br>$F_{(1.0,9.0)}$=0.650; $p$=0.441 | Trial x reward rate<br>$F_{(2.1,18.5)}$=2.71; $p$=0.0915 |

| **Figure 2** | | | |
|---|---|---|---|
| **Panel C– CS-evoked Dopamine** | | | |
| All trials<br>Two-way mixed-effects model | Trial<br>$F_{(5.0,65.1)}$=4.93; $p$=0.0007 | Reward rate<br>$F_{(1.0,13.0)}$=0.000192; $p$=0.989 | Trial x reward rate<br>$F_{(4.3,45.2)}$=1.13; $p$=0.357 |
| Session 1<br>Two-way mixed-effects model | Trial<br>$F_{(2.5,32.3)}$=3.34; **$p$=0.0387** | Reward rate<br>$F_{(1.0,13.0)}$=0.302; $p$=0.592 | Trial x reward rate<br>$F_{(2.3,29.3)}$=4.08; **$p$=0.0234** |
| Session 2<br>Two-way mixed-effects model | Trial<br>$F_{(2.8,36.1)}$=3.21; **$p$=0.0376** | Reward rate<br>$F_{(1.0,13.0)}$=0.232; $p$=0.638 | Trial x reward rate<br>$F_{(1.7,22.0)}$=1.64; $p$=0.219 |
| Session 3<br>Two-way mixed-effects model | Trial<br>$F_{(2.4,26.4)}$=2.31; $p$=0.111 | Reward rate<br>$F_{(1.0,11.0)}$=0.268; $p$=0.615 | Trial x reward rate<br>$F_{(2.8,31.0)}$=0.606; $p$=0.606 |
| Session 4<br>Two-way mixed-effects model | Trial<br>$F_{(2.0,25.7)}$=0.979; $p$=0.388 | Reward rate<br>$F_{(1.0,13.0)}$=0.593; $p$=0.455 | Trial x reward rate<br>$F_{(2.3,29.3)}$=0.251; $p$=0.805 |
| Session 5<br>Two-way mixed-effects model | Trial<br>$F_{(2.7,32.6)}$=0.360; $p$=0.763 | Reward rate<br>$F_{(1.0,12.0)}$=0.233; $p$=0.638 | Trial x reward rate<br>$F_{(2.6,31.0)}$=1.28; $p$=0.297 |
| Session 6<br>Two-way mixed-effects model | Trial<br>$F_{(1.8,23.5)}$=1.71; $p$=0.204 | Reward rate<br>$F_{(1.0,13.0)}$=0.883; $p$=0.365 | Trial x reward rate<br>$F_{(3.0,13.8)}$=1.62; $p$=0.230 |
| **Panel D– US-evoked Dopamine** | | | |
| All trials<br>Two-way mixed-effects model | Trial<br>$F_{(3.8,49.0)}$=9.83; **$p$<0.0001** | Reward rate<br>$F_{(1.0,13.0)}$=2.33; $p$=0.151 | Trial x reward rate<br>$F_{(6.1,64.5)}$=1.04; $p$=0.412 |
| Session 1<br>Two-way mixed-effects model | Trial<br>$F_{(2.0,25.9)}$=1.57; $p$=0.228 | Reward rate<br>$F_{(1.0,13.0)}$=0.0687; $p$=0.797 | Trial x reward rate<br>$F_{(2.4,31.0)}$=2.23; $p$=0.117 |
| Session 2<br>Two-way mixed-effects model | Trial<br>$F_{(2.1,26.9)}$=2.78; $p$=0.0784 | Reward rate<br>$F_{(1.0,13.0)}$=0.932; $p$=0.352 | Trial x reward rate<br>$F_{(2.2,29.1)}$=0.733; $p$=0.504 |
| Session 3<br>Two-way mixed-effects model | Trial<br>$F_{(2.9,32.3)}$=1.54; $p$=0.223 | Reward rate<br>$F_{(1.0,11.0)}$=1.88; $p$=0.198 | Trial x reward rate<br>$F_{(3.2,34.8)}$=0.821; $p$=0.497 |
| Session 4<br>Two-way mixed-effects model | Trial<br>$F_{(2.6,34.0)}$=0.387; $p$=0.736 | Reward rate<br>$F_{(1.0,13.0)}$=1.68; $p$=0.217 | Trial x reward rate<br>$F_{(2.8,36.1)}$=0.845; $p$=0.471 |
| Session 5<br>Two-way mixed-effects model | Trial<br>$F_{(2.3, 27.5)}$=0.447; $p$=0.670 | Reward rate<br>$F_{(1.0,12.0)}$=1.55; $p$=0.237 | Trial x reward rate<br>$F_{(2.2,26.8)}$=0.887; $p$=0.434 |
| Session 6<br>Two-way mixed-effects model | Trial<br>$F_{(2.5,22.3)}$=1.13; $p$=0.350 | Reward rate<br>$F_{(1.0,9.0)}$=3.69; $p$=0.0871 | Trial x reward rate<br>$F_{(1.7,15.4)}$=2.27; $p$=0.142 |
| **Panel E– CS+US Dopamine Response** | | | |
| All trials<br>Two-way mixed-effects model | Trial<br>$F_{(5.1,66.0)}$=3.96; **$p$=.0032** | Reward rate<br>$F_{(1.0,13.0)}$=0.0595; $p$=0.811 | Trial x reward rate<br>$F_{(4.4,46.6)}$=1.21; $p$=0.321 |
| Session 1<br>Two-way mixed-effects model | Trial<br>$F_{(2.5,32.8)}$=4.02; **$p$=0.0201** | Reward rate<br>$F_{(1.0,13.0)}$=0.183; $p$=0.676 | Trial x reward rate<br>$F_{(2.4,31.4)}$=4.49; **$p$=0.0144** |
| Session 2<br>Two-way mixed-effects model | Trial<br>$F_{(2.8,36.7)}$=2.09; $p$=0.122 | Reward rate<br>$F_{(1.0,13.0)}$=0.013; $p$=0.911 | Trial x reward rate<br>$F_{(1.7,22.6)}$=1.67; $p$=0.212 |
| Session 3<br>Two-way mixed-effects model | Trial<br>$F_{(2.4,26.2)}$=2.19; $p$=0.125 | Reward rate<br>$F_{(1.0,11.0)}$=0.583; $p$=0.461 | Trial x reward rate<br>$F_{(2.7,29.8)}$=0.882; $p$=0.452 |
| Session 4<br>Two-way mixed-effects model | Trial<br>$F_{(1.9,25.1)}$=1.31; $p$=0.286 | Reward rate<br>$F_{(1.0,13.0)}$=0.997; $p$=0.336 | Trial x reward rate<br>$F_{(2.2,28.3)}$=0.314; $p$=0.751 |
| Session 5<br>Two-way mixed-effects model | Trial<br>$F_{(2.9,34.6)}$=0.598; $p$=0.615 | Reward rate<br>$F_{(1.0,12.0)}$=0.619; $p$=0.447 | Trial x reward rate<br>$F_{(2.6,31.5)}$=1.04; $p$=0.383 |
| Session 6<br>Two-way mixed-effects model | Trial<br>$F_{(2.0,18.4)}$=2.46; $p$=0.112 | Reward rate<br>$F_{(1.0,9.0)}$=0.918; $p$=0.363 | Trial x reward rate<br>$F_{(2.3,20.8)}$=1.37; $p$=0.277 |

| **Figure 3** | | | |
|---|---|---|---|
| **Panel B – Slope of CS Dopamine Response in NAc Core** | | | |
| All trials<br>Two-way mixed-effects model | Trial<br>$F_{(4.3,39.0)}$=0.865; $p$=0.500 | Reward rate<br>$F_{(1.0,9.0)}$=9.3; **$p$=0.0137** | Trial x reward rate<br>$F_{(4.8,33.0)}$=1.39; $p$=0.256 |
| Session 1<br>Two-way mixed-effects model | Trial<br>$F_{(2.4,14.1)}$=0.616; $p$=0.579 | Reward rate<br>$F_{(1.0,6.0)}$=0.0242; $p$=0.882 | Trial x reward rate<br>$F_{(1.9,11.1)}$=0.251; $p$=0.767 |
| Session 2<br>Two-way mixed-effects model | Trial<br>$F_{(2.7,18.7)}$=0.812; $p$=0.491 | Reward rate<br>$F_{(1.0,7.0)}$=2.83; $p$=0.136 | Trial x reward rate<br>$F_{(2.0,13.8)}$=0.294; $p$=0.747 |
| Session 3<br>Two-way mixed-effects model | Trial<br>$F_{(2.2,19.8)}$=1.26; $p$=0.310 | Reward rate<br>$F_{(1.0,9.0)}$=13.8; **$p$=0.00480** | Trial x reward rate<br>$F_{(2.8,24.9)}$=0.416; $p$=0.712 |
| Session 4<br>Two-way mixed-effects model | Trial<br>$F_{(2.8,22.0)}$=0.920; $p$=0.440 | Reward rate<br>$F_{(1.0,8.0)}$=10.3; **$p$=0.0125** | Trial x reward rate<br>$F_{(2.9,23.1)}$=0.530; $p$=0.660 |

| | | | |
|---|---|---|---|
| Session 5<br>Two-way mixed-effects model | Trial<br>$F_{(1.8,16.3)}$=1.75; $p$=0.206 | Reward rate<br>$F_{(1.0,9.0)}$=4.68; $p$=0.059 | Trial x reward rate<br>$F_{(2.9,26.0)}$=0.841; $p$=0.480 |
| Session 6<br>Two-way mixed-effects model | Trial<br>$F_{(2.8,25.0)}$=1.95; $p$=0.152 | Reward rate<br>$F_{(1.0,9.0)}$=0.0667; $p$=0.802 | Trial x reward rate<br>$F_{(2.8,25.0)}$=0.457; $p$=0.700 |
| **Panel C - Slope of CS Dopamine Response in NAc Shell** | | | |
| All sessions<br>Two-way mixed-effects model | Trial<br>$F_{(4.8,62.2)}$=1.02; $p$=0.410 | Reward rate<br>$F_{(1.0,13.0)}$=0.00573; $p$=0.941 | Trial x reward rate<br>$F_{(4.8,51.1)}$=1.11; $p$=0.364 |
| Session 1<br>Two-way mixed-effects model | Trial<br>$F_{(3.3,43.4)}$=3.10; **$p$=0.0320** | Reward rate<br>$F_{(1.0,13.0)}$=0.381; $p$=0.548 | Trial x reward rate<br>$F_{(2.7,34.8)}$=0.484; $p$=0.675 |
| Session 2<br>Two-way mixed-effects model | Trial<br>$F_{(2.5,32.6)}$=0.884; $p$=0.444 | Reward rate<br>$F_{(1.0,13.0)}$=0.223; $p$=0.645 | Trial x reward rate<br>$F_{(2.5,33.0)}$=1.48; $p$=0.241 |
| Session 3<br>Two-way mixed-effects model | Trial<br>$F_{(2.8,30.5)}$=0.515; $p$=0.661 | Reward rate<br>$F_{(1.0,11.0)}$=0.423; $p$=0.529 | Trial x reward rate<br>$F_{(3.0,33.2)}$=1.42; $p$=0.255 |
| Session 4<br>Two-way mixed-effects model | Trial<br>$F_{(2.6,33.9)}$=1.46; $p$=0.245 | Reward rate<br>$F_{(1.0,13.0)}$=0.133; $p$=0.721 | Trial x reward rate<br>$F_{(2.2,28.7)}$=1.10; $p$=0.352 |
| Session 5<br>Two-way mixed-effects model | Trial<br>$F_{(2.7,32.0)}$=3.40; **$p$=0.0340** | Reward rate<br>$F_{(1.0,12.0)}$=0.0150; $p$=0.905 | Trial x reward rate<br>$F_{(2.9,34.8)}$=2.37; $p$=0.0892 |
| Session 6<br>Two-way mixed-effects model | Trial<br>$F_{(2.5,22.8)}$=0.798; $p$=0.489 | Reward rate<br>$F_{(1.0,9.0)}$=0.0192; $p$=0.893 | Trial x reward rate<br>$F_{(2.4,21.5)}$=2.58; $p$=0.0909 |

| **Figure 4** | | | |
|---|---|---|---|
| **Panel A – CS-evoked Dopamine** | | | |
| All sessions<br>Three-way mixed-effects model | Session<br>$F_{(2.0,43.3)}$=12.2; **$p$<0.0001** | NAc subregion<br>$F_{(1,84)}$=2.08; $p$=0.154 | Reward rate<br>$F_{(1.0,22.0)}$=0.999; $p$=0.331 |
| Session x NAc subregion<br>$F_{(5,84)}$=2.93; **$p$=0.0174** | Session x Reward Rate<br>$F_{(3.4,57.1)}$=1.03; $p$=.393 | NAc subregion x Reward rate<br>$F_{(1,84)}$=0.868; $p$=0.354 | Three-way interaction<br>$F_{(5,84)}$=0.801; $p$=0.552 |
| **Panel B – US-evoked Dopamine** | | | |
| All sessions<br>Three-way mixed-effects model | Session<br>$F_{(1.6,35.2)}$=28.3; **$p$<0.0001** | NAc subregion<br>$F_{(1,84)}$=0.00396; $p$=0.950 | Reward rate<br>$F_{(1.0,22.0)}$=1.41; $p$=0.248 |
| Session x NAc subregion<br>$F_{(5,84)}$=0.557; $p$=0.732 | Session x Reward Rate<br>$F_{(3.6,60.5)}$=0.880; $p$=0.473 | NAc subregion x Reward rate<br>$F_{(1,84)}$=0.0568; $p$=0.812 | Three-way interaction<br>$F_{(5,84)}$=0.721; $p$=0.609 |
| **Panel C – CS+US Dopamine Response** | | | |
| All sessions<br>Three-way mixed-effects model | Session<br>$F_{(2.2,47.8)}$=7.88; **$p$=0.0008** | NAc subregion<br>$F_{(1,84)}$=1.53; $p$=0.219 | Reward rate<br>$F_{(1.0,22.0)}$=3.39; $p$=0.079 |
| Session x NAc subregion<br>$F_{(5,84)}$=3.01; **$p$=0.0152** | Session x Reward Rate<br>$F_{(3.5,58.4)}$=1.34; $p$=0.269 | NAc subregion x Reward rate<br>$F_{(1,84)}$=1.38; $p$=0.243 | Three-way interaction<br>$F_{(5,84)}$=1.07; $p$=0.385 |

| **Figure 5** | | | |
|---|---|---|---|
| **Panel C – Conditioned Responding in Rats with Cannulae in the NAc Core** | | | |
| Session 1-5<br>Three-way mixed-effects model | Trial<br>$F_{(8.8,166.7)}$=13.2; **$p$<0.0001** | Drug<br>$F_{(1,456)}$=6.85; **$p$=0.00910** | Reward rate<br>$F_{(1.0,19.0)}$=1.85; $p$=0.190 |
| Trial x Drug<br>$F_{(24,456)}$=2.29; **$p$=0.0006** | Trial x Reward rate<br>$F_{(2.9,54.8)}$=0.445; $p$=0.715 | Drug x Reward rate<br>$F_{(1,456)}$=0.666; $p$=0.796 | Three-way interaction<br>$F_{(24,456)}$=0.489; $p$=0.982 |
| Session 1<br>Three-way mixed-effects model | Trial<br>$F_{(3.4,64.1)}$=0.874; $p$=0.470 | Drug<br>$F_{(1,76)}$=1.97; $p$=0.165 | Reward rate<br>$F_{(1.0,19.0)}$=1.10; $p$=0.308 |
| Trial x Drug<br>$F_{(4,76)}$=3.28; **$p$=0.0156** | Trial x Reward rate<br>$F_{(2.7,50.5)}$=0.830; $p$=0.471 | Drug x Reward rate<br>$F_{(1,76)}$=0.137; $p$=0.712 | Three-way interaction<br>$F_{(4,76)}$=0.542; $p$=0.705 |
| Session 2<br>Three-way mixed-effects model | Trial<br>$F_{(3.2,61.0)}$=3.13; **$p$=0.0292** | Drug<br>$F_{(1,76)}$=0.878; $p$=0.352 | Reward rate<br>$F_{(1.0,19.0)}$=3.81; $p$=0.0658 |
| Trial x Drug<br>$F_{(4,76)}$=1.40; $p$=0.243 | Trial x Reward rate<br>$F_{(2.3,43.1)}$=0.330; $p$=0.747 | Drug x Reward rate<br>$F_{(1,76)}$=0.114; $p$=0.737 | Three-way interaction<br>$F_{(4,76)}$=0.227; $p$=0.922 |
| Session 3<br>Three-way mixed-effects model | Trial<br>$F_{(3.2,60.1)}$=1.55; $p$=0.210 | Drug<br>$F_{(1,76)}$=4.16; **$p$=0.0449** | Reward rate<br>$F_{(1.0,19.0)}$=1.41; $p$=0.250 |
| Trial x Drug<br>$F_{(4,76)}$=0.208; $p$=0.933 | Trial x Reward rate<br>$F_{(1.8,33.9)}$=0.883; $p$=0.412 | Drug x Reward rate<br>$F_{(1,76)}$=0.0756; $p$=0.784 | Three-way interaction<br>$F_{(4,76)}$=0.198; $p$=0.939 |
| Session 4<br>Three-way mixed-effects model | Trial<br>$F_{(3.2,61.4)}$=4.36; **$p$=0.00630** | Drug<br>$F_{(1,76)}$=3.95; $p$=0.0503 | Reward rate<br>$F_{(1.0,19.0)}$=0.769; $p$=0.391 |
| Trial x Drug<br>$F_{(4,76)}$=0.270; $p$=0.896 | Trial x Reward rate<br>$F_{(1.4,27.0)}$=0.600; $p$=0.501 | Drug x Reward rate<br>$F_{(1,76)}$=0.0683; $p$=0.795 | Three-way interaction<br>$F_{(4,76)}$=0.818; $p$=0.518 |
| Session 5<br>Three-way mixed-effects model | Trial<br>$F_{(3.2,60.1)}$=0.480; $p$=0.708 | Drug<br>$F_{(1,76)}$=10.3; **$p$=0.00190** | Reward rate<br>$F_{(1.0,19.0)}$=0.823; $p$=0.376 |

| | | | |
|---|---|---|---|
| Trial x Drug<br>$F_{(4,76)}=0.322$; $p=0.862$ | Trial x Reward rate<br>$F_{(1.7,33.0)}=0.683$; $p=0.492$ | Drug x Reward rate<br>$F_{(1,76)}=0.257$; $p=0.613$ | Three-way interaction<br>$F_{(4,76)}=1.26$; $p=0.292$ |
| Session 6<br>Three-way mixed-effects model | Trial<br>$F_{(2.9,55.0)}=6.65$; **$p=0.000700$** | Prior drug treatment<br>$F_{(1,76)}=11.5$; **$p=0.00110$** | Reward rate<br>$F_{(1.0,19.0)}=0.787$; $p=0.386$ |
| Trial x Drug<br>$F_{(4,76)}=0.979$; $p=0.424$ | Trial x Reward rate<br>$F_{(1.7,33.2)}=3.83$; **$p=0.0370$** | Prior drug x Reward rate<br>$F_{(1,76)}=0.0862$; $p=0.770$ | Three-way interaction<br>$F_{(4,76)}=0.850$; $p=0.498$ |
| **Panel D – Conditioned Responding in Rats with Cannulae in the NAc Shell** | | | |
| Session 1-5<br>Three-way mixed-effects model | Trial<br>$F_{(6.6,113.0)}=16.6$; **$p<0.0001$** | Drug<br>$F_{(1,408)}=1.16$; $p=0.283$ | Reward rate<br>$F_{(1.0,17.0)}=0.190$; $p=0.669$ |
| Trial x Drug<br>$F_{(24,408)}=1.59$; **$p=0.0395$** | Trial x Reward rate<br>$F_{(3.7,63.1)}=1.09$; $p=0.369$ | Drug x Reward rate<br>$F_{(1,408)}=0.0211$; $p=0.885$ | Three-way interaction<br>$F_{(24,408)}=0.901$; $p=0.602$ |
| Session 1<br>Three-way mixed-effects model | Trial<br>$F_{(3.5,58.9)}=6.89$; **$p=0.0003$** | Drug<br>$F_{(1,68)}=0.0765$; $p=0.783$ | Reward rate<br>$F_{(1.0,17.0)}=0.00428$; $p=0.949$ |
| Trial x Drug<br>$F_{(4,68)}=0.898$; $p=0.470$ | Trial x Reward rate<br>$F_{(3.3,56.5)}=0.387$; $p=0.783$ | Drug x Reward rate<br>$F_{(1,68)}=5.04$; $p=0.0280$ | Three-way interaction<br>$F_{(4,68)}=0.131$; $p=0.970$ |
| Session 2<br>Three-way mixed-effects model | Trial<br>$F_{(3.0,51.3)}=3.03$; **$p=0.0372$** | Drug<br>$F_{(1,68)}=0.221$; $p=0.634$ | Reward rate<br>$F_{(1.0,17.0)}=2.93$; $p=0.105$ |
| Trial x Drug<br>$F_{(4,68)}=0.331$; $p=0.856$ | Trial x Reward rate<br>$F_{(1.8,31.1)}=0.603$; $p=0.540$ | Drug x Reward rate<br>$F_{(1,68)}=0.0125$; $p=0.911$ | Three-way interaction<br>$F_{(4,68)}=0.303$; $p=0.875$ |
| Session 3<br>Three-way mixed-effects model | Trial<br>$F_{(3.4,57.4)}=5.10$; **$p=0.00240$** | Drug<br>$F_{(1,68)}=0.0185$; $p=0.892$ | Reward rate<br>$F_{(1.0,17.0)}=5.84e^{-5}$; $p=0.994$ |
| Trial x Drug<br>$F_{(4,68)}=1.22$; $p=0.309$ | Trial x Reward rate<br>$F_{(2.1,35.6)}=1.88$; $p=0.166$ | Drug x Reward rate<br>$F_{(1,68)}=0.463$; $p=0.499$ | Three-way interaction<br>$F_{(4,68)}=0.816$; $p=0.519$ |
| Session 4<br>Three-way mixed-effects model | Trial<br>$F_{(3.0,50.7)}=5.14$; **$p=0.00360$** | Drug<br>$F_{(1,68)}=0.750$; $p=0.389$ | Reward rate<br>$F_{(1.0,17.0)}=0.194$; $p=0.665$ |
| Trial x Drug<br>$F_{(4,68)}=1.78$; $p=0.144$ | Trial x Reward rate<br>$F_{(1.7,29.3)}=0.648$; $p=0.508$ | Drug x Reward rate<br>$F_{(1,68)}=0.958$; $p=0.331$ | Three-way interaction<br>$F_{(4,68)}=0.538$; $p=0.708$ |
| Session 5<br>Three-way mixed-effects model | Trial<br>$F_{(2.6,44.9)}=0.0669$; $p=0.967$ | Drug<br>$F_{(1,68)}=2.29$; $p=0.135$ | Reward rate<br>$F_{(1.0,17.0)}=0.00926$; $p=0.924$ |
| Trial x Drug<br>$F_{(4,68)}=1.55$; $p=0.197$ | Trial x Reward rate<br>$F_{(1.6,26.9.)}=1.73$; $p=0.201$ | Drug x Reward rate<br>$F_{(1,68)}=0.00292$; $p=0.957$ | Three-way interaction<br>$F_{(4,68)}=0.525$; $p=0.718$ |
| Session 6<br>Three-way mixed-effects model | Trial<br>$F_{(2.5,42.7)}=2.23$; $p=0.108$ | Prior drug treatment<br>$F_{(1,68)}=5.06$; **$p=0.0277$** | Reward rate<br>$F_{(1.0,17.0)}=0.388$; $p=0.542$ |
| Trial x Drug<br>$F_{(4,68)}=0.363$; $p=0.834$ | Trial x Reward rate<br>$F_{(1.9,32.3)}=1.56$; $p=0.225$ | Prior drug x Reward rate<br>$F_{(1,68)}=0.197$; $p=0.659$ | Three-way interaction<br>$F_{(4,68)}=1.73$; $p=0.153$ |

| **Supplementary Figure 1** | | | |
|---|---|---|---|
| **Tray entries during cue** | | | |
| All trials<br>Two-way mixed-effects model | Trial<br>$F_{(29,493)}=4.68$; **$p<0.0001$** | Reward rate<br>$F_{(1,17)}=1.17$; $p=0.295$ | Trial x reward rate<br>$F_{(29,493)}=0.747$; $p=0.829$ |
| Session 1<br>Two-way mixed-effects model | Trial<br>$F_{(4,68)}=6.55$; **$p=0.0002$** | Reward rate<br>$F_{(1,17)}=0.551$; $p=0.468$ | Trial x reward rate<br>$F_{(4,68)}=0.534$; $p=0.711$ |
| Session 2<br>Two-way mixed-effects model | Trial<br>$F_{(4,68)}=4.13$; **$p=0.0047$** | Reward rate<br>$F_{(1,17)}=2.13$; $p=0.163$ | Trial x reward rate<br>$F_{(4,68)}=1.40$; $p=0.244$ |
| Session 3<br>Two-way mixed-effects model | Trial<br>$F_{(4,68)}=1.24$; $p=0.302$ | Reward rate<br>$F_{(1,17)}=2.11$; $p=0.165$ | Trial x reward rate<br>$F_{(4,68)}=0.478$; $p=0.752$ |
| Session 4<br>Two-way mixed-effects model | Trial<br>$F_{(4,68)}=2.45$; $p=0.054$ | Reward rate<br>$F_{(1,17)}=0.148$; $p=0.706$ | Trial x reward rate<br>$F_{(4,68)}=1.07$; $p=0.378$ |
| Session 5<br>Two-way mixed-effects model | Trial<br>$F_{(4,68)}=1.55$; $p=0.197$ | Reward rate<br>$F_{(1,17)}=0.585$; $p=0.455$ | Trial x reward rate<br>$F_{(4,68)}=1.10$; $p=0.363$ |
| Session 6<br>Two-way mixed-effects model | Trial<br>$F_{(4,68)}=0.609$; $p=0.658$ | Reward rate<br>$F_{(1,17)}=0.533$; $p=0.475$ | Trial x reward rate<br>$F_{(4,68)}=0.620$; $p=0.650$ |
| **Tray entries during last 5 s of ITI** | | | |
| All trials<br>Two-way mixed-effects model | Trial<br>$F_{(29,493)}=1.81$; **$p=0.0066$** | Reward rate<br>$F_{(1,17)}=4.68$; **$p=0.045$** | Trial x reward rate<br>$F_{(29,493)}=1.10$; $p=0.337$ |
| Session 1<br>Two-way mixed-effects model | Trial<br>$F_{(4,68)}=0.533$; $p=0.712$ | Reward rate<br>$F_{(1,17)}=0.102$; $p=0.754$ | Trial x reward rate<br>$F_{(4,68)}=0.959$; $p=0.436$ |
| Session 2<br>Two-way mixed-effects model | Trial<br>$F_{(4,68)}=0.843$; $p=0.503$ | Reward rate<br>$F_{(1,17)}=7.08$; **$p=0.0165$** | Trial x reward rate<br>$F_{(4,68)}=1.14$; $p=0.345$ |
| Session 3<br>Two-way mixed-effects model | Trial<br>$F_{(4,68)}=3.62$; **$p=0.0098$** | Reward rate<br>$F_{(1,17)}=0.461$; $p=0.506$ | Trial x reward rate<br>$F_{(4,68)}=0.643$; $p=0.634$ |
| Session 4<br>Two-way mixed-effects model | Trial<br>$F_{(4,68)}=0.949$; $p=0.442$ | Reward rate<br>$F_{(1,17)}=2.65$; $p=0.122$ | Trial x reward rate<br>$F_{(4,68)}=0.684$; $p=0.606$ |

| | | | |
|---|---|---|---|
| Session 5<br>Two-way mixed-effects model | Trial<br>$F_{(4,68)}=2.00$; $p=0.105$ | Reward rate<br>$F_{(1,17)}=0.00702$; $p=0.934$ | Trial x reward rate<br>$F_{(4,68)}=2.02$; $p=0.102$ |
| Session 6<br>Two-way mixed-effects model | Trial<br>$F_{(4,68)}=0.682$; $p=0.607$ | Reward rate<br>$F_{(1,17)}=6.00$; **$p=0.0254$** | Trial x reward rate<br>$F_{(4,68)}=0.695$; $p=0.598$ |

| | | | |
|---|---|---|---|
| **Supplementary Figure 2** | | | |
| **Panel A – Latency to first cue-evoked head entry (animals with electrodes in NAc core)** | | | |
| All trials<br>Two-way mixed-effects model | Trial<br>$F_{(29,493)}=3.78$; **$p<0.0001$** | Reward rate<br>$F_{(1,17)}=0.949$; $p=0.344$ | Trial x reward rate<br>$F_{(29,493)}=0.902$; $p=0.615$ |
| Session 1<br>Two-way mixed-effects model | Trial<br>$F_{(4,68)}=8.39$; **$p<0.0001$** | Reward rate<br>$F_{(1,17)}=0.427$; $p=0.522$ | Trial x reward rate<br>$F_{(4,68)}=1.07$; $p=0.379$ |
| Session 2<br>Two-way mixed-effects model | Trial<br>$F_{(4,68)}=2.70$; **$p=0.0377$** | Reward rate<br>$F_{(1,17)}=0.967$; $p=0.339$ | Trial x reward rate<br>$F_{(4,68)}=0.933$; $p=0.450$ |
| Session 3<br>Two-way mixed-effects model | Trial<br>$F_{(4,68)}=0.772$; $p=0.547$ | Reward rate<br>$F_{(1,17)}=1.13$; $p=0.303$ | Trial x reward rate<br>$F_{(4,68)}=0.365$; $p=0.826$ |
| Session 4<br>Two-way mixed-effects model | Trial<br>$F_{(4,68)}=2.15$; $p=0.085$ | Reward rate<br>$F_{(1,17)}=0.452$; $p=0.511$ | Trial x reward rate<br>$F_{(4,68)}=2.69$; **$p=0.038$** |
| Session 5<br>Two-way mixed-effects model | Trial<br>$F_{(4,68)}=3.49$; **$p=0.0120$** | Reward rate<br>$F_{(1,17)}=0.200$; $p=0.660$ | Trial x reward rate<br>$F_{(4,68)}=1.19$; $p=0.333$ |
| Session 6<br>Two-way mixed-effects model | Trial<br>$F_{(4,68)}=0.506$; $p=0.732$ | Reward rate<br>$F_{(1,17)}=0.959$; $p=0.341$ | Trial x reward rate<br>$F_{(4,68)}=1.04$; $p=0.391$ |

| | | | |
|---|---|---|---|
| **Supplementary Figure 4** | | | |
| **Panel A – Cue-evoked head entries** | | | |
| Session 1<br>Three-way mixed-effects model | Trial<br>$F_{(2.7,50.8)}=5.93$; **$p=0.00220$** | Drug<br>$F_{(1,76)}=3.39$; $p=0.0696$ | Reward rate<br>$F_{(1.0,19.0)}=0.380$; $p=0.545$ |
| Trial x Drug<br>$F_{(4,76)}=1.40$; $p=0.242$ | Trial x Reward rate<br>$F_{(2.4,46.0)}=0.468$; $p=0.666$ | Drug x Reward rate<br>$F_{(1,76)}=0.789$; $p=0.377$ | Three-way interaction<br>$F_{(4,76)}=0.582$; $p=0.677$ |
| Session 2<br>Three-way mixed-effects model | Trial<br>$F_{(3.2,61.5)}=14.8$; **$p<0.0001$** | Drug<br>$F_{(1,76)}=3.88$; $p=0.0526$ | Reward rate<br>$F_{(1.0,19.0)}=2.45$; $p=0.134$ |
| Trial x Drug<br>$F_{(4,76)}=1.67$; $p=0.165$ | Trial x Reward rate<br>$F_{(2.0,38.4)}=0.495$; $p=0.615$ | Drug x Reward rate<br>$F_{(1,76)}=0.992$; $p=0.322$ | Three-way interaction<br>$F_{(4,76)}=0.335$; $p=0.854$ |
| Session 3<br>Three-way mixed-effects model | Trial<br>$F_{(3.3,63.4)}=5.26$; **$p=0.00190$** | Drug<br>$F_{(1,76)}=8.53$; **$p=0.00460$** | Reward rate<br>$F_{(1.0,19.0)}=2.30$; $p=0.146$ |
| Trial x Drug<br>$F_{(4,76)}=1.78$; $p=0.142$ | Trial x Reward rate<br>$F_{(1.7,33.2)}=1.26$; $p=0.294$ | Drug x Reward rate<br>$F_{(1,76)}=0.377$; $p=0.541$ | Three-way interaction<br>$F_{(4,76)}=0.166$; $p=0.955$ |
| Session 4<br>Three-way mixed-effects model | Trial<br>$F_{(3.2,61.7)}=9.21$; **$p<0.0001$** | Drug<br>$F_{(1,76)}=3.77$; $p=0.0558$ | Reward rate<br>$F_{(1.0,19.0)}=2.22$; $p=0.153$ |
| Trial x Drug<br>$F_{(4,76)}=1.22$; $p=0.308$ | Trial x Reward rate<br>$F_{(1.2,23.6)}=1.54$; $p=0.231$ | Drug x Reward rate<br>$F_{(1,76)}=0.0887$; $p=0.767$ | Three-way interaction<br>$F_{(4,76)}=0.214$; $p=0.930$ |
| Session 5<br>Three-way mixed-effects model | Trial<br>$F_{(3.3,62.0)}=3.03$; **$p=0.0322$** | Drug<br>$F_{(1,76)}=10.2$; **$p=0.00210$** | Reward rate<br>$F_{(1.0,19.0)}=0.790$; $p=0.385$ |
| Trial x Drug<br>$F_{(4,76)}=0.792$; $p=0.534$ | Trial x Reward rate<br>$F_{(1.5.28.7)}=3.07$; $p=0.0745$ | Drug x Reward rate<br>$F_{(1,76)}=0.260$; $p=0.612$ | Three-way interaction<br>$F_{(4,76)}=1.12$; $p=0.352$ |
| Session 6<br>Three-way mixed-effects model | Trial<br>$F_{(3.0,57.1)}=12.9$; **$p<0.0001$** | Prior drug treatment<br>$F_{(1,76)}=7.93$; **$p=0.00620$** | Reward rate<br>$F_{(1.0,19.0)}=1.79$; $p=0.196$ |
| Trial x Drug<br>$F_{(4,76)}=0.721$; $p=0.580$ | Trial x Reward rate<br>$F_{(1.6,31.3)}=2.08$; $p=0.149$ | Prior drug x Reward rate<br>$F_{(1,76)}=0.359$; $p=0.551$ | Three-way interaction<br>$F_{(4,76)}=1.29$; $p=0.281$ |
| **Panel B – ITI head entries** | | | |
| Session 1<br>Three-way mixed-effects model | Trial<br>$F_{(2.6,49.4)}=16.9$; **$p<0.0001$** | Drug<br>$F_{(1,76)}=1.45$; $p=0.232$ | Reward rate<br>$F_{(1.0,19.0)}=67.4$; **$p<0.0001$** |
| Trial x Drug<br>$F_{(4,76)}=0.561$; $p=0.692$ | Trial x Reward rate<br>$F_{(1.4,25.8)}=15.3$; **$p=0.0002$** | Drug x Reward rate<br>$F_{(1,76)}=1.55$; $p=0.218$ | Three-way interaction<br>$F_{(4,76)}=0.457$; $p=0.767$ |
| Session 2<br>Three-way mixed-effects model | Trial<br>$F_{(3.3,63.5)}=16.0$; **$p<0.0001$** | Drug<br>$F_{(1,76)}=0.706$; $p=0.403$ | Reward rate<br>$F_{(1.0,19.0)}=116$; **$p<0.0001$** |
| Trial x Drug<br>$F_{(4,76)}=0.966$; $p=0.431$ | Trial x Reward rate<br>$F_{(1.5,29.2)}=12.8$; **$p=0.0003$** | Drug x Reward rate<br>$F_{(1,76)}=1.62$; $p=0.207$ | Three-way interaction<br>$F_{(4,76)}=1.78$; $p=0.142$ |
| Session 3<br>Three-way mixed-effects model | Trial<br>$F_{(3.1,58.0)}=4.94$; **$p=0.00380$** | Drug<br>$F_{(1,76)}=1.83$; $p=0.180$ | Reward rate<br>$F_{(1.0,19.0)}=157$; **$p<0.0001$** |
| Trial x Drug<br>$F_{(4,76)}=1.23$; $p=0.304$ | Trial x Reward rate<br>$F_{(1.9,36.6)}=3.76$; **$p=0.0341$** | Drug x Reward rate<br>$F_{(1,76)}=3.20$; $p=0.0776$ | Three-way interaction<br>$F_{(4,76)}=0.949$; $p=0.440$ |
| Session 4<br>Three-way mixed-effects model | Trial<br>$F_{(3.0,56.1)}=2.17$; $p=0.103$ | Drug<br>$F_{(1,76)}=0.0206$; $p=0.886$ | Reward rate<br>$F_{(1.0,19.0)}=108$; **$p<0.0001$** |

| | | | |
|---|---|---|---|
| Trial x Drug<br>$F_{(4,76)}=1.17$; $p=0.330$ | Trial x Reward rate<br>$F_{(1.5,28.0)}=1.36$; $p=0.266$ | Drug x Reward rate<br>$F_{(1,76)}=0.822$; $p=0.367$ | Three-way interaction<br>$F_{(4,76)}=1.49$; $p=0.214$ |
| Session 5<br>Three-way mixed-effects model | Trial<br>$F_{(3.5,66.5)}=8.44$; **$p<0.0001$** | Drug<br>$F_{(1,76)}=1.40$; $p=0.241$ | Reward rate<br>$F_{(1.0,19.0)}=69.8$; **$p<0.0001$** |
| Trial x Drug<br>$F_{(4,76)}=0.894$; $p=0.472$ | Trial x Reward rate<br>$F_{(1.6,30.3)}=6.26$; **$p=0.00850$** | Drug x Reward rate<br>$F_{(1,76)}=2.47$; $p=0.120$ | Three-way interaction<br>$F_{(4,76)}=0.512$; $p=0.727$ |
| Session 6<br>Three-way mixed-effects model | Trial<br>$F_{(2.5,47.0)}=10.2$; **$p<0.0001$** | Prior drug treatment<br>$F_{(1,76)}=6.28e^{-6}$; $p=0.998$ | Reward rate<br>$F_{(1.0,19.0)}=107$; **$p<0.0001$** |
| Trial x Drug<br>$F_{(4,76)}=1.89$; $p=0.120$ | Trial x Reward rate<br>$F_{(1.6,29.5)}=5.06$; **$p=0.0190$** | Prior drug x Reward rate<br>$F_{(1,76)}=0.0856$; $p=0.771$ | Three-way interaction<br>$F_{(4,76)}=2.41$; $p=0.0563$ |
| **Panel C – Latency to first cue-evoked head entry** | | | |
| Session 1<br>Three-way mixed-effects model | Trial<br>$F_{(3.3,62.4)}=13.9$; **$p<0.0001$** | Drug<br>$F_{(1,76)}=0.760$; $p=0.386$ | Reward rate<br>$F_{(1.0,19.0)}=6.25$; **$p=0.0218$** |
| Trial x Drug<br>$F_{(4,76)}=0.639$; $p=0.639$ | Trial x Reward rate<br>$F_{(3.1,59.5)}=0.474$; $p=0.710$ | Drug x Reward rate<br>$F_{(1,76)}=0.830$; $p=0.365$ | Three-way interaction<br>$F_{(4,76)}=1.04$; $p=0.392$ |
| Session 2<br>Three-way mixed-effects model | Trial<br>$F_{(3.5,67.4)}=16.8$; **$p<0.0001$** | Drug<br>$F_{(1,76)}=0.826$; $p=0.366$ | Reward rate<br>$F_{(1.0,19.0)}=0.607$; $p=0.446$ |
| Trial x Drug<br>$F_{(4,76)}=0.418$; $p=0.795$ | Trial x Reward rate<br>$F_{(2.4,45.2)}=0.261$; $p=0.808$ | Drug x Reward rate<br>$F_{(1,76)}=0.535$; $p=0.467$ | Three-way interaction<br>$F_{(4,76)}=0.0873$; $p=0.986$ |
| Session 3<br>Three-way mixed-effects model | Trial<br>$F_{(3.3,62.0)}=7.57$; **$p=0.0001$** | Drug<br>$F_{(1,76)}=2.11$; $p=0.150$ | Reward rate<br>$F_{(1.0,19.0)}=2.57$; $p=0.126$ |
| Trial x Drug<br>$F_{(4,76)}=0.597$; $p=0.666$ | Trial x Reward rate<br>$F_{(2.8,53.9)}=1.37$; $p=0.263$ | Drug x Reward rate<br>$F_{(1,76)}=1.67$; $p=0.201$ | Three-way interaction<br>$F_{(4,76)}=0.631$; $p=0.642$ |
| Session 4<br>Three-way mixed-effects model | Trial<br>$F_{(3.5,65.9)}=11.9$; **$p<0.0001$** | Drug<br>$F_{(1,76)}=0.120$; $p=0.731$ | Reward rate<br>$F_{(1.0,19.0)}=1.65$; $p=0.215$ |
| Trial x Drug<br>$F_{(4,76)}=0.183$; $p=0.947$ | Trial x Reward rate<br>$F_{(1.6,31.1)}=0.769$; $p=0.448$ | Drug x Reward rate<br>$F_{(1,76)}=0.271$; $p=0.605$ | Three-way interaction<br>$F_{(4,76)}=0.752$; $p=0.560$ |
| Session 5<br>Three-way mixed-effects model | Trial<br>$F_{(3.1,57.9)}=3.97$; **$p=0.0118$** | Drug<br>$F_{(1,76)}=0.369$; $p=0.546$ | Reward rate<br>$F_{(1.0,19.0)}=1.05$; $p=0.318$ |
| Trial x Drug<br>$F_{(4,76)}=0.412$; $p=0.800$ | Trial x Reward rate<br>$F_{(2.1,39.6)}=5.70$; **$p=0.00610$** | Drug x Reward rate<br>$F_{(1,76)}=0.700$; $p=0.407$ | Three-way interaction<br>$F_{(4,76)}=0.786$; $p=0.538$ |
| Session 6<br>Three-way mixed-effects model | Trial<br>$F_{(3.3,62.6)}=19.7$; **$p<0.0001$** | Prior drug treatment<br>$F_{(1,76)}=1.76$; $p=0.189$ | Reward rate<br>$F_{(1.0,19.0)}=2.74$; $p=0.114$ |
| Trial x Drug<br>$F_{(4,76)}=0.104$; $p=0.981$ | Trial x Reward rate<br>$F_{(2.9,55.2)}=3.85$; **$p=0.0151$** | Prior drug x Reward rate<br>$F_{(1,76)}=0.342$; $p=0.560$ | Three-way interaction<br>$F_{(4,76)}=1.60$; $p=0.183$ |

| **Supplementary Figure 5** | | | |
|---|---|---|---|
| **Panel A – Cue-evoked head entries** | | | |
| Session 1<br>Three-way mixed-effects model | Trial<br>$F_{(2.9,49.4)}=23.0$; **$p<0.0001$** | Drug<br>$F_{(1,68)}=0.285$; $p=0.595$ | Reward rate<br>$F_{(1.0,17.0)}=0.243$; $p=0.628$ |
| Trial x Drug<br>$F_{(4,68)}=2.09$; $p=0.0915$ | Trial x Reward rate<br>$F_{(2.0,34.7)}=2.89$; $p=0.0680$ | Drug x Reward rate<br>$F_{(1,68)}=4.51$; **$p=0.0373$** | Three-way interaction<br>$F_{(4,68)}=0.971$; $p=0.429$ |
| Session 2<br>Three-way mixed-effects model | Trial<br>$F_{(3.0,51.2)}=6.05$; **$p<0.00130$** | Drug<br>$F_{(1,68)}=0.191$; $p=0.663$ | Reward rate<br>$F_{(1.0,17.0)}=2.96$; $p=0.103$ |
| Trial x Drug<br>$F_{(4,68)}=0.173$; $p=0.951$ | Trial x Reward rate<br>$F_{(1.6,27.4)}=0.786$; $p=0.440$ | Drug x Reward rate<br>$F_{(1,68)}=0.000364$; $p=0.985$ | Three-way interaction<br>$F_{(4,68)}=0.831$; $p=0.510$ |
| Session 3<br>Three-way mixed-effects model | Trial<br>$F_{(3.2,53.8)}=9.84$; **$p<0.0001$** | Drug<br>$F_{(1,68)}=0.0302$; $p=0.863$ | Reward rate<br>$F_{(1.0,17.0)}=0.968$; $p=0.339$ |
| Trial x Drug<br>$F_{(4,68)}=0.610$; $p=0.657$ | Trial x Reward rate<br>$F_{(1.7,29.7)}=1.84$; $p=0.180$ | Drug x Reward rate<br>$F_{(1,68)}=0.0631$; $p=0.803$ | Three-way interaction<br>$F_{(4,68)}=1.19$; $p=0.323$ |
| Session 4<br>Three-way mixed-effects model | Trial<br>$F_{(2.7,46.2)}=7.18$; **$p=0.0007$** | Drug<br>$F_{(1,68)}=0.211$; $p=0.648$ | Reward rate<br>$F_{(1.0,17.0)}=1.01$; $p=0.328$ |
| Trial x Drug<br>$F_{(4,68)}=0.507$; $p=0.731$ | Trial x Reward rate<br>$F_{(1.6,26.7)}=0.378$; $p=0.638$ | Drug x Reward rate<br>$F_{(1,68)}=0.852$; $p=0.359$ | Three-way interaction<br>$F_{(4,68)}=0.348$; $p=0.845$ |
| Session 5<br>Three-way mixed-effects model | Trial<br>$F_{(2.7,45.5)}=0.120$; $p=0.933$ | Drug<br>$F_{(1,68)}=2.70$; $p=0.105$ | Reward rate<br>$F_{(1.0,17.0)}=0.228$; $p=0.639$ |
| Trial x Drug<br>$F_{(4,68)}=0.707$; $p=0.590$ | Trial x Reward rate<br>$F_{(1.6,27.8)}=1.79$; $p=0.190$ | Drug x Reward rate<br>$F_{(1,68)}=0.00195$; $p=0.965$ | Three-way interaction<br>$F_{(4,68)}=0.573$; $p=0.684$ |
| Session 6<br>Three-way mixed-effects model | Trial<br>$F_{(3.3,55.5)}=4.13$; **$p<0.00860$** | Prior drug treatment<br>$F_{(1,68)}=3.97$; $p=0.0503$ | Reward rate<br>$F_{(1.0,17.0)}=0.644$; $p=0.434$ |
| Trial x Drug<br>$F_{(4,68)}=0.504$; $p=0.733$ | Trial x Reward rate<br>$F_{(1.8,29.9)}=1.94$; $p=0.166$ | Prior drug x Reward rate<br>$F_{(1,68)}=0.112$; $p=0.740$ | Three-way interaction<br>$F_{(4,68)}=1.28$; $p=0.285$ |
| **Panel B – ITI head entries** | | | |

| | | | |
|---|---|---|---|
| Session 1<br>Three-way mixed-effects model | Trial<br>$F_{(3.3,55.3)}$=9.84; ***p*<0.0001** | Drug<br>$F_{(1,68)}$=$4.90e^{-5}$; *p*=0.994 | Reward rate<br>$F_{(1.0,17.0)}$=99.2; ***p*<0.0001** |
| Trial x Drug<br>$F_{(4,68)}$=1.97; *p*=0.108 | Trial x Reward rate<br>$F_{(1.8,29.8)}$=10.7; ***p*=0.0005** | Drug x Reward rate<br>$F_{(1,68)}$=0.00429; *p*=0.948 | Three-way interaction<br>$F_{(4,68)}$=1.63; *p*=0.177 |
| Session 2<br>Three-way mixed-effects model | Trial<br>$F_{(3.5,59.1)}$=6.54; ***p*=0.0004** | Drug<br>$F_{(1,68)}$=2.33; *p*=0.132 | Reward rate<br>$F_{(1.0,17.0)}$=97.2; ***p*<0.0001** |
| Trial x Drug<br>$F_{(4,68)}$=0.670; *p*=0.615 | Trial x Reward rate<br>$F_{(1.4,23.6)}$=2.24; *p*=0.142 | Drug x Reward rate<br>$F_{(1,68)}$=2.85; *p*=0.0962 | Three-way interaction<br>$F_{(4,68)}$=0.961; *p*=0.435 |
| Session 3<br>Three-way mixed-effects model | Trial<br>$F_{(2.7,46.1)}$=3.68; ***p*=0.0218** | Drug<br>$F_{(1,68)}$=0.264; *p*=0.609 | Reward rate<br>$F_{(1.0,17.0)}$=98.4; ***p*<0.0001** |
| Trial x Drug<br>$F_{(4,68)}$=0.683; *p*=0.606 | Trial x Reward rate<br>$F_{(2.2,36.8)}$=3.64; ***p*=0.0331** | Drug x Reward rate<br>$F_{(1,68)}$=1.01; *p*=0.318 | Three-way interaction<br>$F_{(4,68)}$=1.10; *p*=0.363 |
| Session 4<br>Three-way mixed-effects model | Trial<br>$F_{(2.8,47.3)}$=1.97; *p*=0.136 | Drug<br>$F_{(1,68)}$=0.00156; *p*=0.969 | Reward rate<br>$F_{(1.0,17.0)}$=79.2; ***p*<0.0001** |
| Trial x Drug<br>$F_{(4,68)}$=0.533; *p*=0.712 | Trial x Reward rate<br>$F_{(1.6,28.0)}$=0.941; *p*=0.356 | Drug x Reward rate<br>$F_{(1,68)}$=0.0257; *p*=0.873 | Three-way interaction<br>$F_{(4,68)}$=0.288; *p*=0.885 |
| Session 5<br>Three-way mixed-effects model | Trial<br>$F_{(3.1,52.6)}$=0.817; *p*=0.494 | Drug<br>$F_{(1,68)}$=0.0482; *p*=0.827 | Reward rate<br>$F_{(1.0,17.0)}$=38.8; ***p*<0.0001** |
| Trial x Drug<br>$F_{(4,68)}$=2.99; ***p*=0.0246** | Trial x Reward rate<br>$F_{(1.1,19.5)}$=1.51; *p*=0.238 | Drug x Reward rate<br>$F_{(1,68)}$=$2.35e^{-5}$; *p*=0.996 | Three-way interaction<br>$F_{(4,68)}$=0.909; *p*=0.464 |
| Session 6<br>Three-way mixed-effects model | Trial<br>$F_{(3.0,51.8)}$=2.50; *p*=0.0687 | Prior drug treatment<br>$F_{(1,68)}$=0.00485; *p*=0.945 | Reward rate<br>$F_{(1.0,17.0)}$=63.6; ***p*<0.0001** |
| Trial x Drug<br>$F_{(4,68)}$=0.858; *p*=0.494 | Trial x Reward rate<br>$F_{(1.5,24.8)}$=1.89; *p*=0.179 | Prior drug x Reward rate<br>$F_{(1,68)}$=0.0203; *p*=0.887 | Three-way interaction<br>$F_{(4,68)}$=0.312; *p*=0.869 |
| **Panel C – Latency to first cue-evoked head entry** | | | |
| Session 1<br>Three-way mixed-effects model | Trial<br>$F_{(3.5,60.1)}$=18.7; ***p*<0.0001** | Drug<br>$F_{(1,68)}$=0.918; *p*=0.341 | Reward rate<br>$F_{(1.0,17.0)}$=2.79; *p*=0.113 |
| Trial x Drug<br>$F_{(4,68)}$=1.60; *p*=0.185 | Trial x Reward rate<br>$F_{(2.6,44.5)}$=1.12; *p*=0.346 | Drug x Reward rate<br>$F_{(1,68)}$=1.52; *p*=0.222 | Three-way interaction<br>$F_{(4,68)}$=0.837; *p*=0.506 |
| Session 2<br>Three-way mixed-effects model | Trial<br>$F_{(3.3,56.2)}$=5.44; ***p*=0.0017** | Drug<br>$F_{(1,68)}$=0.749; *p*=0.390 | Reward rate<br>$F_{(1.0,17.0)}$=4.23; *p*=0.0554 |
| Trial x Drug<br>$F_{(4,68)}$=1.53; *p*=0.203 | Trial x Reward rate<br>$F_{(2.3,39.2)}$=0.933; *p*=0.413 | Drug x Reward rate<br>$F_{(1,68)}$=0.0311; *p*=0.861 | Three-way interaction<br>$F_{(4,68)}$=0.499; *p*=0.737 |
| Session 3<br>Three-way mixed-effects model | Trial<br>$F_{(3.4,57.4)}$=10.5; ***p*<0.0001** | Drug<br>$F_{(1,68)}$=0.00158; *p*=0.968 | Reward rate<br>$F_{(1.0,17.0)}$=1.41; *p*=0.252 |
| Trial x Drug<br>$F_{(4,68)}$=0.657; *p*=0.624 | Trial x Reward rate<br>$F_{(2.7,46.4)}$=0.945; *p*=0.420 | Drug x Reward rate<br>$F_{(1,68)}$=0.0404; *p*=0.841 | Three-way interaction<br>$F_{(4,68)}$=0.501; *p*=0.735 |
| Session 4<br>Three-way mixed-effects model | Trial<br>$F_{(3.2,54.1)}$=4.80; ***p*=0.00420** | Drug<br>$F_{(1,68)}$=0.462; *p*=0.499 | Reward rate<br>$F_{(1.0,17.0)}$=2.69; *p*=0.120 |
| Trial x Drug<br>$F_{(4,68)}$=0.249; *p*=0.909 | Trial x Reward rate<br>$F_{(2.5,41.8)}$=0.875; *p*=0.444 | Drug x Reward rate<br>$F_{(1,68)}$=1.03; *p*=0.313 | Three-way interaction<br>$F_{(4,68)}$=0.116; *p*=0.976 |
| Session 5<br>Three-way mixed-effects model | Trial<br>$F_{(3.1,52.6)}$=1.10; *p*=0.357 | Drug<br>$F_{(1,68)}$=1.23; *p*=0.272 | Reward rate<br>$F_{(1.0,17.0)}$=0.358; *p*=0.558 |
| Trial x Drug<br>$F_{(4,68)}$=0.621; *p*=0.649 | Trial x Reward rate<br>$F_{(1.9,31.6.)}$=2.08; *p*=0.145 | Drug x Reward rate<br>$F_{(1,68)}$=0.197; *p*=0.659 | Three-way interaction<br>$F_{(4,68)}$=1.59; *p*=0.188 |
| Session 6<br>Three-way mixed-effects model | Trial<br>$F_{(3.5,58.7)}$=12.3; ***p*<0.0001** | Prior drug treatment<br>$F_{(1,68)}$=6.23; ***p*=0.0150** | Reward rate<br>$F_{(1.0,17.0)}$=2.80; *p*=0.113 |
| Trial x Drug<br>$F_{(4,68)}$=0.297; *p*=0.879 | Trial x Reward rate<br>$F_{(2.2,38.0)}$=2.74; *p*=0.0716 | Prior drug x Reward rate<br>$F_{(1,68)}$=0.0326; *p*=0.857 | Three-way interaction<br>$F_{(4,68)}$=1.52; *p*=0.207 |
